# Clustering of host N-glycans licenses *Toxoplasma* rhoptry discharge

**DOI:** 10.1101/2025.10.16.682961

**Authors:** Dylan Valleau, Mudita Goyal, Justin M. Roberts, Frances Male, Gary Ward, Sebastian Lourido

## Abstract

Apicomplexan parasites must discharge the contents of specialized organelles called rhoptries into host cells to initiate the process of invasion. This process requires the prior recognition and binding of the host cell by proteins released from another set of parasite organelles, the micronemes. However, the host-parasite interactions required for rhoptry discharge are largely unknown. Here we performed a host-cell directed genome-wide screen for host factors required for rhoptry discharge from *Toxoplasma gondii,* the causative agent of toxoplasmosis. The screen identified host N-glycosylation and cholesterol biosynthesis as pathways required for normal rhoptry discharge. A trimeric microneme complex, MIC1/4/6, interfaces with both pathways by binding host N-glycans to cluster proteins in a process dependent on host plasma membrane cholesterol. The process can be inhibited by depletion of host cholesterol or competition with exogenous glycans. This clustering of host factors by MIC1/4/6 likely prepares the host membrane for rhoptry discharge, delineating a new step in the *Toxoplasma* invasion process.

## INTRODUCTION

Invasion of host cells is required to establish the replicative niche of apicomplexan parasites. Studies of zoites from multiple stages of *Toxoplasma gondii*, *Plasmodium* spp., and *Cryptosporidium* spp. have identified common steps in invasion including initial attachment, reorientation, penetration (or enveloping in the case of *Cryptosporidium*) and fission of the parasitophorous vacuole from the host plasma membrane ^1–3^. In *T. gondii*, the entire process is completed in less than a minute ^4^ progression through the sequence of events is highly-regulated and depends on the discharge of micronemes and rhoptries, specialized organelles that secrete proteins from the apical end of parasites ^5,6^. Micronemes are numerous and their contents are released in waves to support egress, gliding motility, and invasion ^7^. By contrast, zoites have limited numbers of the larger club-shaped rhoptries, ranging from a single one in *Cryptosporidium* spp. sporozoites to up to a dozen in *Toxoplasma* tachyzoites. The timing of rhoptry discharge is precisely regulated to deliver the arsenal of effector proteins directly into host cells that then drives invasion and host remodeling ^5,8,9^. The prior secretion of certain microneme proteins is necessary, but not sufficient, for rhoptry discharge ^10–13^.

Microneme protein secretion initiates egress and continues throughout the extracellular phase of the parasite life cycle, up to the point of invasion. Micronemes fuse with the apical plasma membrane in response to waves of Ca^2+^ release from both intracellular stores and the extracellular environment ^7,14^. The signaling pathways that regulate microneme fusion include cGMP production and PKG upstream of the Ca^2+^ surge ^15^, and calcium-dependent protein kinases (CDPKs), notably CDPK1, in the transduction of the signals ^5,6^. DOC2 and ferlins have also been implicated in the apical fusion of micronemes ^16,17^, which releases soluble luminal contents into the extracellular space and incorporates transmembrane proteins and associated complexes onto the parasite plasma membrane. Microneme proteins then perform several functions critical for invasion, including mediating adhesion to the host during gliding motility and licensing rhoptry discharge upon host cell contact ^8,18^.

Several processes contribute to the overall efficiency of rhoptry discharge. Gliding motility and attachment are not strictly required but enhance rates of rhoptry discharge by promoting proper apposition of the parasite apical end and the host cell membrane. Indeed, the effects of gliding motility can be excluded in rhoptry discharge assays by blocking actin polymerization (e.g. Cytochalasin D or Mycalolide B) or disrupting key motors (e.g. MyoH) ^19^. Analogously, the formation of the moving junction increases parasite attachment, which can be interpreted as increased rhoptry discharge when visually quantifying the release of rhoptry contents in proximity to attached parasites (i.e. evacuole formation) ^20^; however, loss of moving junction formation does not affect rhoptry discharge when the process is evaluated directly ^10,21^. It should be noted that current methods for measuring rhoptry discharge do not formally distinguish between the release of rhoptry proteins from the parasite and their translocation into host cells, so discharge will be used as a unified term for the ensemble process. For example, the microneme-secreted adhesins MIC2 and AMA1 participate in parasite adhesion and moving junction formation, respectively ^22–24^, but their disruption does not affect rhoptry discharge when the process is measured while blocking gliding motility ^10,11^. By contrast, several microneme proteins have been specifically linked to rhoptry discharge in *T. gondii* without generally affecting secretion of other microneme proteins, disrupting gliding motility, or interfering with organelle biogenesis; such factors include MIC7, MIC8, and components of the CLAMP and CRMP complexes ^10–12,25^. The CLAMP and CRMP complexes are the most conserved of these factors as their core proteins are generally encoded in all apicomplexan genomes; however, their precise function in rhoptry discharge remains unknown.

Contrasting with the membrane fusion of micronemes, rhoptry discharge occurs via an elaborate rhoptry secretion apparatus (RSA). Ultrastructural studies have shown that the RSA is constitutively embedded in the plasma membrane at the apical tip of the parasite, visualized as a rosette in *T. gondii* ^26^. The rhoptries are docked to this apparatus via an apical vesicle (AV), so content release is thought to occur through two regulated steps: fusion of the rhoptry with the AV and AV fusion with the parasite plasma membrane at the RSA site^8,26–28^. Recent high-speed imaging and electrophysiology has uncovered that rhoptry discharge by *T. gondii* is associated with the transient perforation of the host cell plasma membrane ^21,29^. The erythrocytic blood-stages of *P. falciparum* also form a pore in the erythrocyte plasma membrane prior to invasion, suggesting permeabilization is a common feature of invasion for other apicomplexans, though the identity of the perforating factors remains unclear in both species ^1,30^.

Disruption of the host membrane provides a plausible mechanism for translocation of rhoptry proteins directly into the host cytoplasm, following discharge through the RSA. Translocation of *T. gondii* rhoptry effectors can be used to measure rhoptry discharge directly or indirectly. Discharge of the rhoptry protein ROP16 leads to the phosphorylation and nuclear accumulation of the host transcription factor STAT6, which can readily be quantified by immunofluorescence ^31^. Alternatively, fusion proteins with the rhoptry protein Toxofilin may be used to translocate the Cre recombinase or a beta-lactamase, enabling indirect detection of rhoptry discharge by measuring the activity of fused enzymes ^32–34^. Notably, measurement of beta-lactamase activity in host cells remains the most rapid and scalable method to quantify rhoptry discharge. After translocation, some rhoptry proteins, including the rhoptry neck (RON) complex, are embedded into the host plasma membrane, allowing the C-terminal tail of RON2 to act as an anchor for the microneme protein AMA1, which in turn is tethered to the parasite plasma membrane and the parasite actomyosin system for force generation ^18,20,24^. Together, this RON2-AMA1 interaction forms the core of the moving junction, through which the parasite invades.

Several host factors contribute to invasion. *T. gondii* adhesion can be disrupted by blocking interactions between sulfated proteoglycans like heparin and MIC2 ^22,35–37^. Host cholesterol also appears to be required for efficient invasion, but the interacting parasite factors remain unknown ^38,39^. In *Plasmodium falciparum*, the clade-specific *Pf*RCR complex mediates binding to host basigin and α2,6-linked Neu5Ac (sialic acid) on red blood cells, and is thought to be involved in rhoptry discharge ^9,40,41^. Many of the *T. gondii* rhoptry discharge factors noted above have adhesive domains that similarly suggest affinity for specific host cell ligands, although, notably, no specific host receptors have been identified.

We set out to identify the host factors specifically required for *T. gondii* rhoptry discharge, employing a host-directed genome-wide CRISPR/Cas9-based screen. The analysis revealed that host cholesterol and N-glycosylation are critical for rhoptry discharge. Orthogonal to known rhoptry discharge factors, we further identify the microneme complex that intersects with the two pathways to license rhoptry discharge and promote infection.

## RESULTS

### A genome-wide host knockout CRISPR screen reveals host determinants of rhoptry discharge

To identify host determinants of *T. gondii* rhoptry discharge, we developed a screening approach using a previously established FRET-based assay for rhoptry discharge ^34^. In this assay, rhoptry discharge is directly assessed from parasites expressing the secreted rhoptry effector Toxofilin (Txfn) conjugated to beta-lactamase (βla). After reporter discharge into mammalian host cells, they can be loaded with the coumarin-fluorescein FRET substrate CCF4-AM. Initially CCF4 fluoresces green (520 nm) when excited at a wavelength of 405 nm. Txfn-βla then cleaves the β-lactam ring in CCF4, uncoupling the FRET and resulting in a shift to blue fluorescence (447 nm; **Figure 1A**). To exclude any confounding effects from parasite invasion, we introduced Txfn-βla into a *T. gondii* strain engineered for conditional knockdown of MyoH, a parasite myosin required for motility and invasion but dispensable for microneme and rhoptry discharge ^19,42^. Conditional knockdown of MyoH was achieved using a rapamycin-dimerizable Cre recombinase (DiCre) that causes post-transcriptional regulation of MyoH levels using the U1 system (DiCre/MyoH cKD/Txfn-βla) ^43,44^.

**Figure 1.**
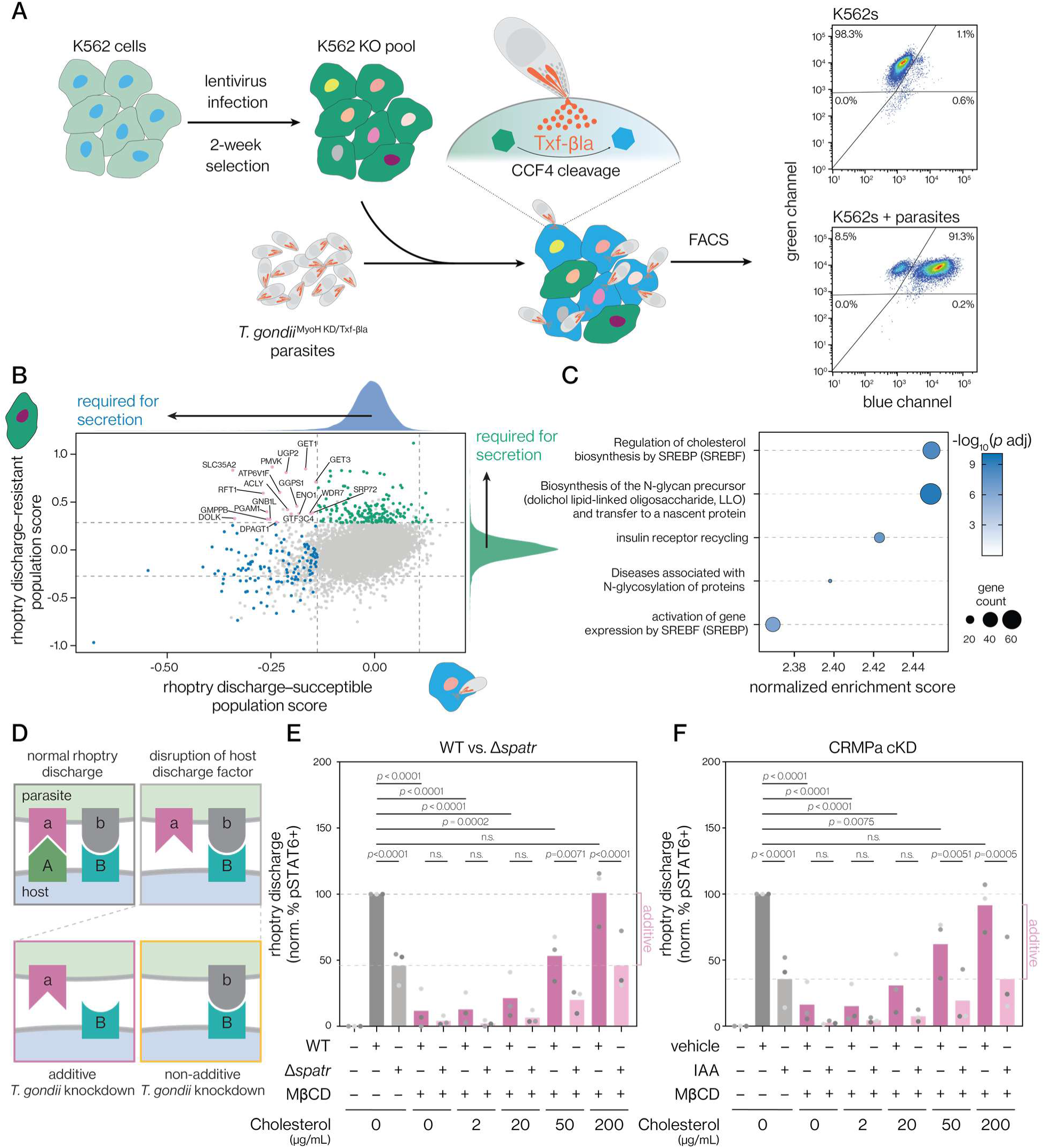
A host-directed screen reveals genes required for rhoptry discharge. **(A)** Design of the rhoptry discharge screen. A genome-wide knockout pool was generated in the human leukemia K562 cell line using CRISPR-based gene disruption. The host cell pool was challenged with rapamycin-treated DiCre/MyoH cKD/Txfn-βla *T. gondii* parasites, then loaded with CCF4-AM and sorted into populations refractory or susceptible to rhoptry discharge. Sample plots show the relative signal of CCF4 with or without parasite-delivered Txfn-βla. **(B)** MAGeCK MLE analysis of guide RNA abundance in sorted host cell populations. Beta-score calculated by the MLE pipeline are shown. The distribution of scores for each population is displayed along each axis. Genes are colored based on the enrichment of targeting guide RNAs in the population refractory to rhoptry discharge (green), depletion from the susceptible population (blue), or presence in both groups (pink). Inclusion in each group was based on s.d > 3 and Wald test *p-*value < 0.05. Dashed lines represent the s.d. equivalent to 3 for each distribution. **(C)** GSEA of the top human Reactome pathways required for rhoptry discharge into host cells. **(D)** Scheme to determine whether host and parasite factors participate in common or parallel interaction pathways. An additive effect is expected from host and parasite perturbations disrupting different pathways, whereas non-additive interactions result from them disrupting a common pathway. **(E–F)** Cholesterol was depleted from human fibroblasts with MβCD and then reconstituted to different levels. Rhoptry discharge was evaluated based on the relative percent of pSTAT6– positive nuclei after incubation with Δ*spatr* or wild-type parental parasites (**E**) or a CRMPa conditional knockdown strain treated with vehicle or IAA (**F**). Mean of *n* = 3 biological replicates is plotted; *p*-values are derived from two-way ANOVA testing with multiple comparison FDR correction using the Benjamini, Krieger, and Yekutieli two-stage step-up method.

Human erythroleukemia K562 cells were transduced with a pooled 5-guide (97,888 guides) lentivirus CRISPR-based library targeting the entire human genome for knockout (19,665 protein-coding genes) ^45^. On days 13 and 14 post transduction, the pools of K562 knockout host cells were allowed to interact with DiCre/MyoH cKD/Txfn-βla parasites that had been treated with a 2-hour rapamycin treatment 2 days prior to induce MyoH knockdown. Parasites were added at a multiplicity of infection (MOI) of 20, which had been optimized to achieve 90% CCF4 conversion among the K562 pool (**Figure 1A**). After 1 hour of interaction, CCF4-AM was added and K562 cells were sorted into two populations: green (refractory to rhoptry discharge) and blue (susceptible to rhoptry discharge). We performed the screen in two independently transduced K562 pools, each infected on two separate days, for a total of 4 experimental replicates. Integrated guide RNAs were amplified from each population and their relative abundances in each sample were measured by next-generation sequencing.

Using the maximum-likelihood estimation (MLE) MAGeCK pipeline, we compared sorted populations to their respective unsorted controls (**Figure 1B**). 200 factors were deemed critical for rhoptry discharge based on either their significant enrichment in the discharge-resistant (green) population or their significant depletion from the discharge-sensitive (blue) populations, with 125 factors overlapping between the two categories, and 18 being both greater than 3 s.d. from the mean in both populations and significant in at least one population. Notably, proteins localized to the plasma membrane were largely absent among putative rhoptry discharge factors (**Figure S1A**) ^46–48^, consistent with *T. gondii* not depending on any single receptor protein and explaining the parasite’s broad tropism. However, several top scoring genes included components of complexes involved in membrane protein biogenesis or trafficking, including many components of the GET, EMC, and SRP complexes (**Figure 1B**), revealing that the display of surface proteins is generally important for rhoptry discharge.

To categorize host genes important for rhoptry discharge, we combined the scores from refractory and susceptible populations, and performed gene set enrichment analysis (GSEA) against the Reactome database ^49^. Pathways related to N-glycan processing and biosynthesis, and cholesterol biosynthesis and regulation appeared essential for rhoptry discharge (**Figure 1C**, **S1B–C**). By contrast, pathways previously found to be involved in parasite attachment— notably, the biosynthesis of heparin sulfate, which is bound by the adhesion-related microneme protein MIC2—are not strongly enriched for putative discharge factors (**Figure S1D**), consistent with our screen specifically targeting the step of rhoptry discharge ^35,37^.

Investigating the genes required for rhoptry discharge within the identified cholesterol pathways finds both cholesterol-specific synthesis genes like MSMO1 ^50^ as well as many mevalonate pathway genes (e.g. MVK and PMVK), which supply prenoid precursors for cholesterol biosynthesis, geranylation, and farnesylation (**Figure S1C**). The transcriptional regulators of sterol production, SREBP/SERBF, as well as some of their downstream genes (e.g. FASN, HMGCR) also appeared to be important for rhoptry discharge ^51^.

In the N-glycan pathway, a striking number of the genes responsible for making the essential dolichol glycan precursors and shuttling them to the ER lumen were detected, including DOLK, DPAGT1, and RFT1, as well as genes required for synthesizing and trimming the core N-glycan structure, such as many ALG genes (**Figure S1B**) ^52^. The requirement for synthesis of N-linked glycoproteins, which are predominantly displayed on the host cell surface, is consistent with the involvement of the aforementioned EMC and SRP complexes in rhoptry discharge. Interestingly, some of the rhoptry discharge determinants, such as NUS1, are important for synthesis of the dolichol N-glycan precursor from mevalonate-pathway products, suggesting a possible integration of these two categories of rhoptry discharge factors ^51,53^.

### Host plasma membrane cholesterol is required for rhoptry discharge independently of known parasite factors

Host cell cholesterol has previously been found to be important for parasite replication and invasion ^38,39,54,55^. Depletion of cholesterol from host cells blocks formation of evacuoles, a well-established marker of rhoptry discharge ^39,56^. Cholesterol affects fluidity of membranes and is required for various processes that involve membrane reshaping such as pore formation and protein clustering ^57^. Host cholesterol accumulates at the site of invasion, but dissipates quickly from the nascent parasitophorous vacuole ^54^. This is consistent with host membrane cholesterol being required for the invasion process, although its direct relationship to rhoptry discharge has not been fully evaluated.

In *T. gondii* there are two microneme protein complexes required for normal rhoptry discharge and conserved throughout apicomplexans: the CRMP and CLAMP complexes ^10–12^. As cholesterol is present in all mammalian cells, we sought to determine whether these essential complexes depend on host cholesterol for their roles in rhoptry discharge. We reasoned that interactions between host and parasite factors could be evaluated by examining how parasites deficient in specific rhoptry discharge factors (e.g. CLAMP or CRMP complex knockdowns) respond to the simultaneous disruption of candidate host factors. In this approach, simultaneous inhibition of host and parasite factors that act on the same pathway should inhibit rhoptry discharge to the same extent as inhibiting either component alone, resulting in a non-additive experimental readout. In contrast, if the dually targeted parasite and host factors play independent roles in rhoptry discharge, we should see a more severe additive effect, as we would be disrupting multiple host-parasite interactions simultaneously (**Figure 1D**).

To disrupt the CLAMP and CRMP complexes, we used a SPATR knockout (Δ*spatr*) and an auxin-inducible CRMPa knockdown (CRMPa-mAID), respectively ^10–12,58^. The cyclodextrin MβCD was used to deplete host cells of cholesterol ^39,59^, which can then be reconstituted into the plasma membrane at various concentrations to exclude non-specific or permanent effects from cholesterol depletion (e.g. changes in cell signaling or viability). Parasites were treated with mycalolide B, an irreversible inhibitor of actin polymerization that blocks motility and invasion without affecting rhoptry discharge, analogously to MyoH knockdown ^10^. Rhoptry discharge was assessed by monitoring the fraction of host cells displaying phosphorylation and nuclear translocation of the transcription factor STAT6, which is induced by the secreted rhoptry effector ROP16 ^31^. This alternative measure of rhoptry discharge avoids possible changes in the loading efficiency of CCF4-AM following cholesterol depletion or other host membrane perturbations. Rhoptry discharge was strongly inhibited by host cell cholesterol depletion, and completely rescued at the highest levels of cholesterol reconstitution (**Figure 1E-F**), confirming the dependency of rhoptry discharge specifically on host plasma membrane cholesterol. However, we observed additive effects between the CRMP and CLAMP complex perturbations and host membrane cholesterol depletion, revealing that these host and parasite perturbations affect parallel host-parasite interactions pathways that independently contribute to rhoptry discharge.

### *T. gondii* rhoptry discharge is dependent on host cell galactose in N-glycans

N-glycosylation was the other major pathway implicated in rhoptry discharge by our screen. Two of the top four host genes with the strongest effects on rhoptry discharge, SLC35A2 and UGP2, are required for galactosylation and glucosylation of N-glycans, respectively ^60–63^. SLC35A2 transports UDP-Galactose (UDP-Gal) into the Golgi, where it provides the monosaccharide to growing glycan chains. UGP2 is a metabolic enzyme involved in synthesis of UDP-Glucose (UDP-Glc) (**Figure 2A**). UDP-Glc is also the main source of UDP-Gal under standard glucose-rich cell culture conditions through interconversion of the two precursors by the enzyme GALE (**Figure 2A**) ^62–64^. We generated knockouts of UGP2 and SLC35A2 in K562 cells, and confirmed that both mutant cell lines showed robust staining with GS-II lectin, which binds terminal GlcNAc moieties exposed on glycans when capping galactosylation is not performed (**Figure 2B**) ^61^. As expected, due to GALE-mediated interconversion growth for three days in the presence of galactose partially restored galactosylation in UGP2 KO but not SLC35A2 KO cells, since cells lacking the latter transporter cannot import UDP-Gal into the Golgi despite its increased synthesis from galactose supplementation (**Figure 2B**). As predicted by the screen, loss of either UGP2 or SLC35A2 significantly reduced rhoptry discharge (**Figure 2C**). Moreover, rhoptry discharge into UGP2 KO host cells could be partially rescued by growth of the cell line in galactose, but no such rescue was observed for SLC35A2 KO (**Figure 2D**). These experiments indicate that import of UDP-Gal into the Golgi is critical for host cell susceptibility to rhoptry discharge.

**Figure 2.**
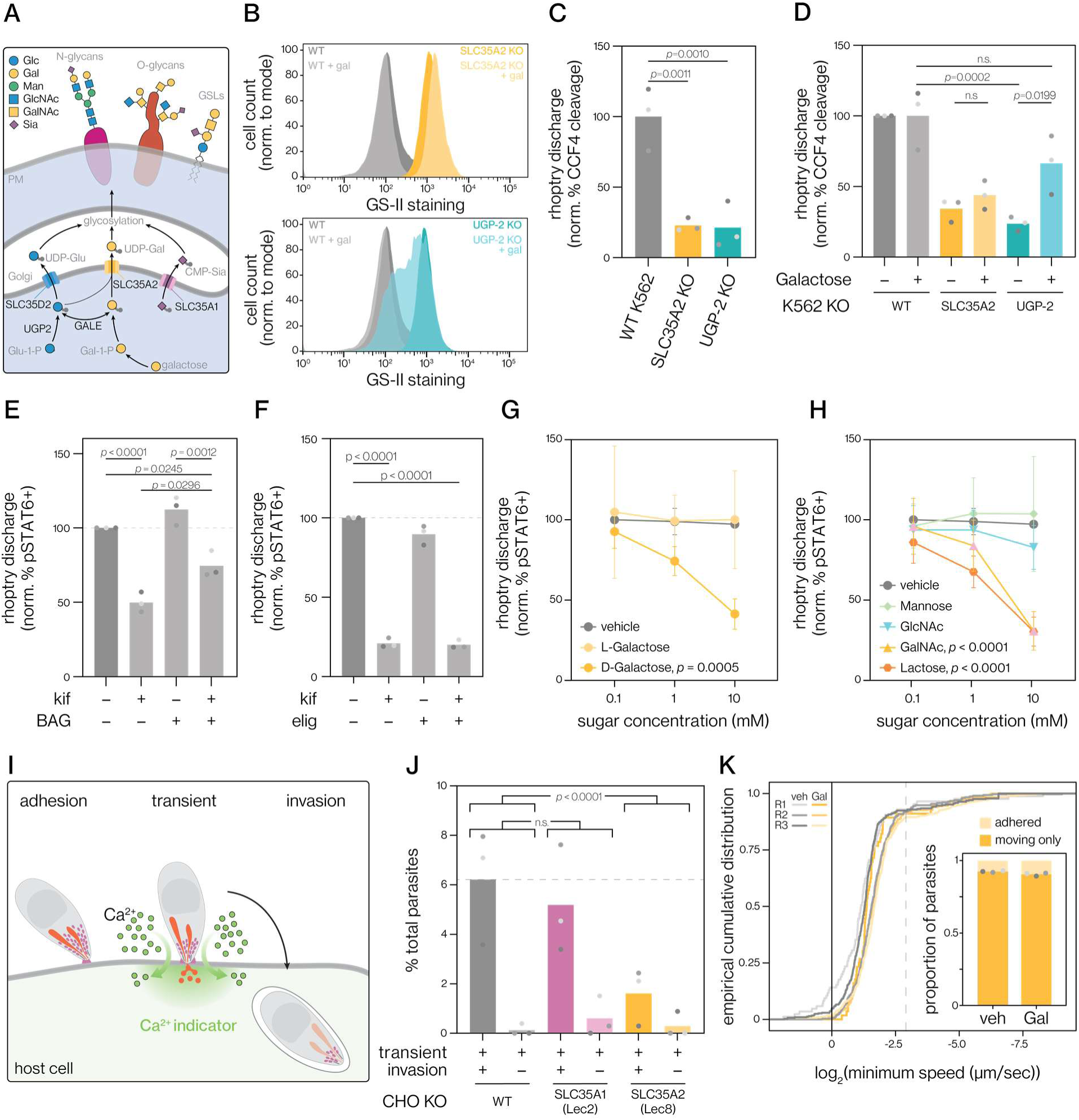
Galactosylation of host N-glycans is required for rhoptry discharge. **(A)** Simplified schematic of synthesis, transport, and utilization of UDP-Glc, UDP-Gal, and CMP-Sia precursors in various mammalian glycosylation pathways. **(B)** Flow cytometry of K562 cell lines in which UGP-2 or SLC35A2 have been knocked out (KO). Cells were labeled with GS-II, which recognizes uncapped surface GlcNAc. Galactose supplementation (25 mM, 3 days) rescues surface glycosylation patterns for UGP-2 KO but not SLC35A2 KO cells. **(C)** Rhoptry discharge was assessed in the KO cell lines as the percentage of host cells with βla-cleaved CCF4 relative to WT host cells, following challenge with rapamycin-treated DiCre/MyoH cKD/Txfn-βla parasites. Mean of *n* = 3 biological replicates is plotted; *p*-values derived from one-way ANOVA with Fisher’s LSD post-hoc test. **(D)** Rhoptry discharge into host KO cell lines grown in presence of vehicle or galactose (25 mM, 3 days) was assessed as the percentage of host cells with βla-cleaved CCF4 relative to vehicle-treated WT K562 cells, following challenge with rapamycin-treated DiCre/MyoH cKD/Txfn-βla parasites. Mean of *n* = 3 biological replicates is plotted; *p*-values derived from one-way ANOVA with Tukey’s post hoc test. **(E-F)** Human fibroblasts were treated with the N-glycosylation inhibitor kif (10 μM) alone or in combination with the O-glycosylation inhibitor BAG (2 mM) for 3 days (**E**), or the glycosphingolipid inhibitor elig (2 μM) for 5 days (**F**). Host cells were then challenged with wild-type parasites and rhoptry discharge was assessed by the relative percent of pSTAT6–positive nuclei. Mean of *n* = 3 biological replicates is plotted; *p*-values derived from one-way ANOVA with Tukey’s post hoc test. **(G-H)** Rhoptry discharge in the presence of varying amounts of galactose stereoisomers (**G**) or galactose-containing glycans (**H**). Rhoptry discharge was assessed by the relative percent of pSTAT6-positive nuclei after challenge with wild-type parasites. Each replicate was normalized to the vehicle control of the 0.1 mM condition; *p*-values for the comparison to the vehicle control at 10 mM concentration are provided when significant, derived from two-way ANOVA testing with multiple comparison FDR correction using the Benjamini, Krieger, and Yekutieli two-stage step-up method. Mean of *n* = 3 biological replicates is plotted with error bars indicating s.d. Data from **G** and **H** were analyzed together in the same statistical model. **(I)** Measurement of host cell transient perforation and parasite invasion. In a microfluidic microscopy slide, parasites adhere to host cells loaded with the Ca^2+^ indicator Cal-520. Upon rhoptry discharge, transient host cell perforation allows Ca^2+^ influx into the host cytosol, increasing Cal-520 fluorescence. Parasites are then individually tracked to identify those that invade subsequently to the permeabilization event. **(J)** The perforation and invasion of CHO cells lacking galactosylation (SLC35A2 KO) or sialylation (SLC35A1 KO) by wild-type parasites. Mean of *n* = 3 biological replicates is plotted; *p*-values are derived from Fisher’s exact test. **(K)** The impact of galactose on parasite adhesion to host cells was assessed by flowing wild-type parasites over human fibroblasts grown in microfluidic chambers. The cumulative distribution of the log_2_ transformed minimum speed of each parasite for each replicate is plotted. The log2(minimum speed) threshold used to classify parasites as moving and adhered is displayed (dashed grey line), and the relative ratio of moving and adherent parasites is displayed in the inset. Mean of *n* = 3 biological replicates is plotted.

Galactose is present in many types of glycans ^52^; however, N-glycosylation was the dominant glycosylation pathway identified in our screen based on GSEA and enrichment of N-glycan specific genes like DOLK, DPAGT1, RFT1, DDOST, and several ALG genes (**Figure 1C**, **S1B**). By contrast, enzymes specific to O-glycosylation and glycosphingolipid synthesis were largely absent among the genes identified as rhoptry discharge factors. To confirm whether N-glycans are the primary surface-exposed glycans involved in rhoptry discharge we employed selective inhibitors: kifunensine (kif), an inhibitor of alpha-mannosidases required for formation of complex and hybrid N-glycans ^65,66^; and benzyl-α-GalNAc (BAG), a competitive inhibitor of mucin-type O-glycan initiation ^67,68^. After treating replicating human fibroblasts for three days with kif, we saw significant inhibition of rhoptry discharge (**Figure 2E**). By contrast, treatment with BAG had no effect on rhoptry discharge. Interestingly, a combination of both BAG and kif led to a moderate rescue of rhoptry discharge compared to kif alone, consistent with the notion that compensation between N- and O-glycosylation pathways can dictate the resulting balance of cell glycosylation ^69–71^. Another major class of surface-exposed glycosylation, glycosphingolipids, can be depleted by inhibiting the rate-limiting enzyme in their synthesis, UGCG, with eliglustat (elig) ^72,73^. Treatment of dividing human fibroblasts with elig for five days lowered glycosphingolipid levels based on surface binding of fluorescently-conjugated Cholera toxin subunit B (CTB), which binds to the glycosphingolipid ganglioside GM1 (**Figure S2A**). Elig treatment had a negligible effect on rhoptry discharge, compared to kif treatment for the same period of time, further demonstrating the importance of N-glycans (**Figure 2F**). Taken together, these results point to galactose-containing N-glycans as the primary surface glycans required for normal rhoptry discharge.

Many pathogenic bacteria, viruses, and parasites bind host glycans ^74^ and these interactions can be competitively disrupted by the addition of exogenous glycans in culture or *in vivo* ^75^. *T. gondii* adhesion and invasion can be reduced by incubation with heparan and sialic acid, respectively ^10,37,76,77^. We therefore tested whether exogenous galactose could competitively disrupt rhoptry discharge. When added together with parasites—i.e. without prior incubation—D-galactose significantly inhibited rhoptry discharge, in a dose-dependent manner (**Figure 2G**). By contrast, the enantiomer L-galactose had no effect, excluding non-specific effects on the osmolarity or composition of the media. Structurally related sugars like GalNAc and the galactose-containing disaccharide lactose similarly inhibited rhoptry discharge, while GlcNAc and mannose had no effect (**Figure 2H**). Collectively, these results suggest that exogenous galactose and related sugars can competitively inhibit parasite binding of surface-exposed galactose-containing host N-glycans.

### Host N-glycosylation is required for rhoptry discharge independently of the CLAMP and CRMP complexes

To further study the relationship between glycosylation and rhoptry discharge, we turned to Chinese Hamster Ovary (CHO) cells, for which characterized mutants are available disrupting N-glycosylation maturation via MGAT1 (Lec1), CMP-sialic acid Golgi transport via SLC35A1 (Lec2), and UDP-Gal Golgi transport via SLC35A2 (Lec8) ^78,79^. Using the lectins that recognize mannose in N-glycans (ConA), GlcNAc and Neu5Ac (WGA), and terminal GlcNAc (GS-II), we confirmed these cell lines had the expected surface glycosylation patterns (**Figure S2B-C**), providing a set of adherent cell lines in a second species to probe the role of host glycosylation in *T. gondii* rhoptry discharge.

As an alternative approach to examining rhoptry discharge, we examined transient host cell permeabilization during *T. gondii* infection. Recent work has demonstrated that parasites transiently perforate the host cell membrane prior to invasion in a manner that depends on rhoptry discharge factors like the CLAMP complex ^21,29^. To examine which stage of invasion is affected by disruption of host galactosylation, we measured host cell perforation by imaging Ca^2+^ transients in host cells during invasion (**Figure 2I**) ^21^. Consistent with a role for host galactosylation in the initiation of rhoptry discharge, we observed fewer permeabilization events in SLC35A2 KO cells, and this coincided with fewer invasion events. By contrast, SLC35A1 KO host cells, deficient in sialylation, were perforated and invaded at similar rates to wild-type (**Figure 2J**). Like the Txfn-βla and pSTAT6 assays, host cell perforation requires rhoptry discharge ^21^, but perforation likely precedes the protein translocation required for a positive Txfn-βla and pSTAT6 signal. The reduction in the frequency of perforation is therefore consistent with host galactosylation being required early in the process of rhoptry discharge.

Because rhoptry discharge occurs downstream of attachment, we used a flow-based adhesion assay to evaluate this initial interaction with host cells. In this assay, parasites are flowed over a confluent monolayer of host cells and imaged continuously to capture the frequency and duration of attachment events. The results demonstrated that exogenous galactose did not alter the general ability of parasites to adhere to human fibroblasts (**Figure 2K**), despite its effect on the subsequent invasion step of rhoptry discharge (**Figure 2G–H**).

The effect of exogenous galactose—interfering with rhoptry discharge but not adhesion—is mirrored by the knockdown of rhoptry discharge factors like the CRMP and CLAMP complexes ^10–12^, motivating the evaluation of additivity between these host and parasite pathways. Using a genetic interaction approach as with cholesterol depletion above, we examined the interaction between galactosyl sugars and the CRMP and CLAMP complexes. Although exogenous galactose and lactose robustly impaired rhoptry discharge into human fibroblasts, we observed that this effect was additive with the disruption of either the CLAMP complex (Δ*spatr*) or CRMP complex (CRMPa cKD; **Figure 3A–B**). A similar additive interaction could be observed between global disruption of N-glycosylation and Δ*spatr* parasites (**Figure S3A**). As expected, mannose had no effect on rhoptry discharge. However, exogenous sialic acid (Neu5Ac) modestly inhibited rhoptry discharge, suggesting that host sialic acid may also marginally contribute to attachment or rhoptry discharge independently of the the CRMP or CLAMP complexes (**Figure 3A–B**).

**Figure 3.**
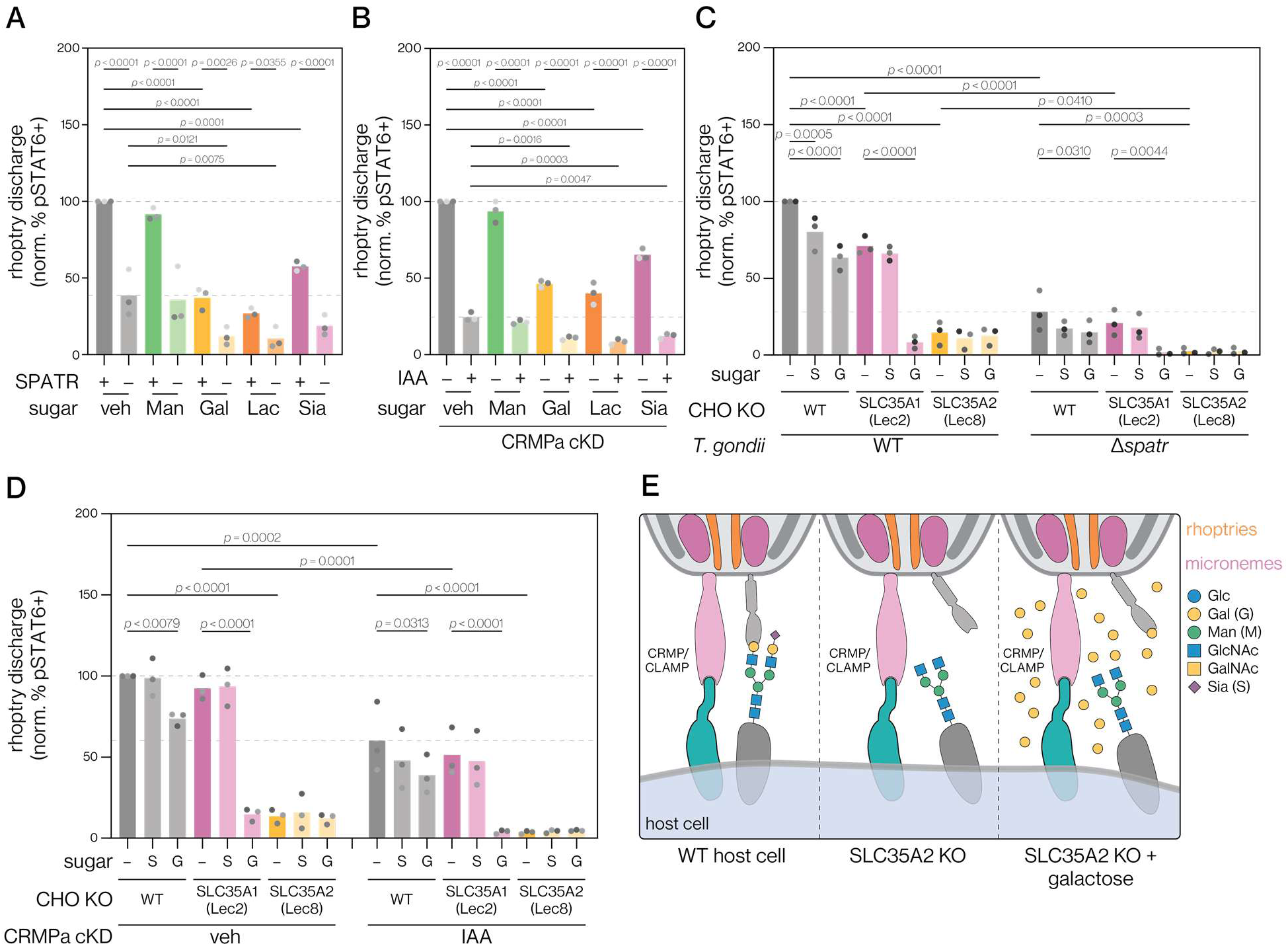
The CRMP and CLAMP complexes are not involved in binding galactosyl N-glycans during rhoptry discharge. **(A-B)** Individual and combined effects of exogenous sugars and genetic disruption of the CLAMP complex (**A**) or the CRMP complex (**B**). 10 mM of the indicated sugar was added with the parasites (Man, mannose; Gal, galactose; Lac, lactose; Sia, sialic acid). Human fibroblasts were incubated with wild-type (SPATR +) or Δ*spatr* parasites (SPATR –) (**A**), or CRMPa cKD parasites grown in presence of IAA or vehicle control (**B**). Rhoptry discharge was assessed by the relative percent of pSTAT6–positive nuclei. Mean of *n* = 3 biological replicates is plotted; *p*-values for each experiment derived from two-way ANOVA with Tukey’s post hoc multiple comparison test. **(C-D)** Individual or combined effects of genetic perturbation of host and parasite pathways together with exogenous sugars. Wild-type CHO cells or mutants in glycosylation pathways (SLC35A1 and SLC35A2 KOs) were challenged with parasites disrupted in CLAMP (**C**) or CRMP (**D**) complexes, as above. 10 mM of each indicated exogenous sugar was added with the parasites (G, galactose; S, sialic acid). Rhoptry discharge was assessed by the relative percent of pSTAT6–positive nuclei. Mean of *n* = 3 biological replicates is plotted; *p*-values are derived from two-way ANOVA testing with multiple comparison FDR correction using the Benjamini, Krieger, and Yekutieli two-stage step-up method. **(E)** Model explaining the non-additive effect of combining exogenous galactose and SLC35A2 KO, implying that the effect of host galactosylation is independent of the pathways regulated by CLAMP and CRMP complexes.

To confirm the additive interactions between host galactosylation and the parasite complexes, we performed pSTAT6 assays in the CHO knockout cells, combined with exogenously added glycans. This confirmed that the impaired rhoptry discharge into SLC35A2 KO cells could not be further inhibited by exogenous galactose, although it was inhibited by disruption of the CLAMP or CRMP complexes (**Figure 3C–D**). Interestingly, galactose and sialic acid suppressed rhoptry discharge more modestly in CHO cells compared to human fibroblasts, suggesting that the balance of factors influencing rhoptry discharge may differ between cell types or host species. Nevertheless, in SLC35A1 KO cells, which lack capping sialic acids and therefore have complex N-glycans predominantly terminating in galactose, exogenous galactose addition strongly blocked rhoptry discharge, while sialic acid had no added effect, as expected (**Figure 3C–D**). As with human fibroblasts, mannose had no effect on rhoptry discharge into CHO cells (**Figure S3B-C**). Taken together, these results suggest there are other parasite factors responsible for recognizing galactose-containing N-glycans in rhoptry discharge (**Figure 3E**).

### The MIC1/4/6 complex interacts with galactose-containing glycans

Our results suggest that the CRMP and CLAMP complexes are not responsible for the recognition of galactose-containing N-glycans on the host cell. The other known rhoptry-discharge factor, MIC8, lacks detectable glycan-binding domains ^13^. Therefore, we sought to identify parasite proteins capable of binding galactose-containing glycans. Inspired by previous approaches to discover lactose-binding parasite proteins ^80–82^, we flowed *T. gondii* lysate over agarose beads with conjugated galactose, lactose, or mannose moieties. After stringent washing, we specifically eluted galactose, lactose, or mannose-binding proteins by flowing high concentrations (>100 mM) of the cognate sugars over their respective resins and used mass spectrometry to identify bound parasite proteins. As mannose is not involved in rhoptry discharge (**Figure 2H, S3B-C**), this condition serves as a negative control. MIC1, MIC4, MIC6 were specifically enriched by lactose and galactose beads compared to mannose (**Figure 4A**). This is consistent with previous studies that identified MIC1, MIC4 and MIC6 form a trimeric complex, and that MIC1 and MIC4 can be isolated by lactose bead pull down ^81,83^. MIC4 is known to be a terminal galactose-binding lectin, likely responsible for facilitating the binding to lactose and galactose resins for the MIC1/4/6 complex ^77^. The next most abundant protein was MIC13, which is a paralog of MIC1 ^76^. No other microneme proteins were significantly enriched, suggesting the MIC1/4/6 complex may be the dominant galactose-interacting microneme complex in *T. gondii*.

**Figure 4.**
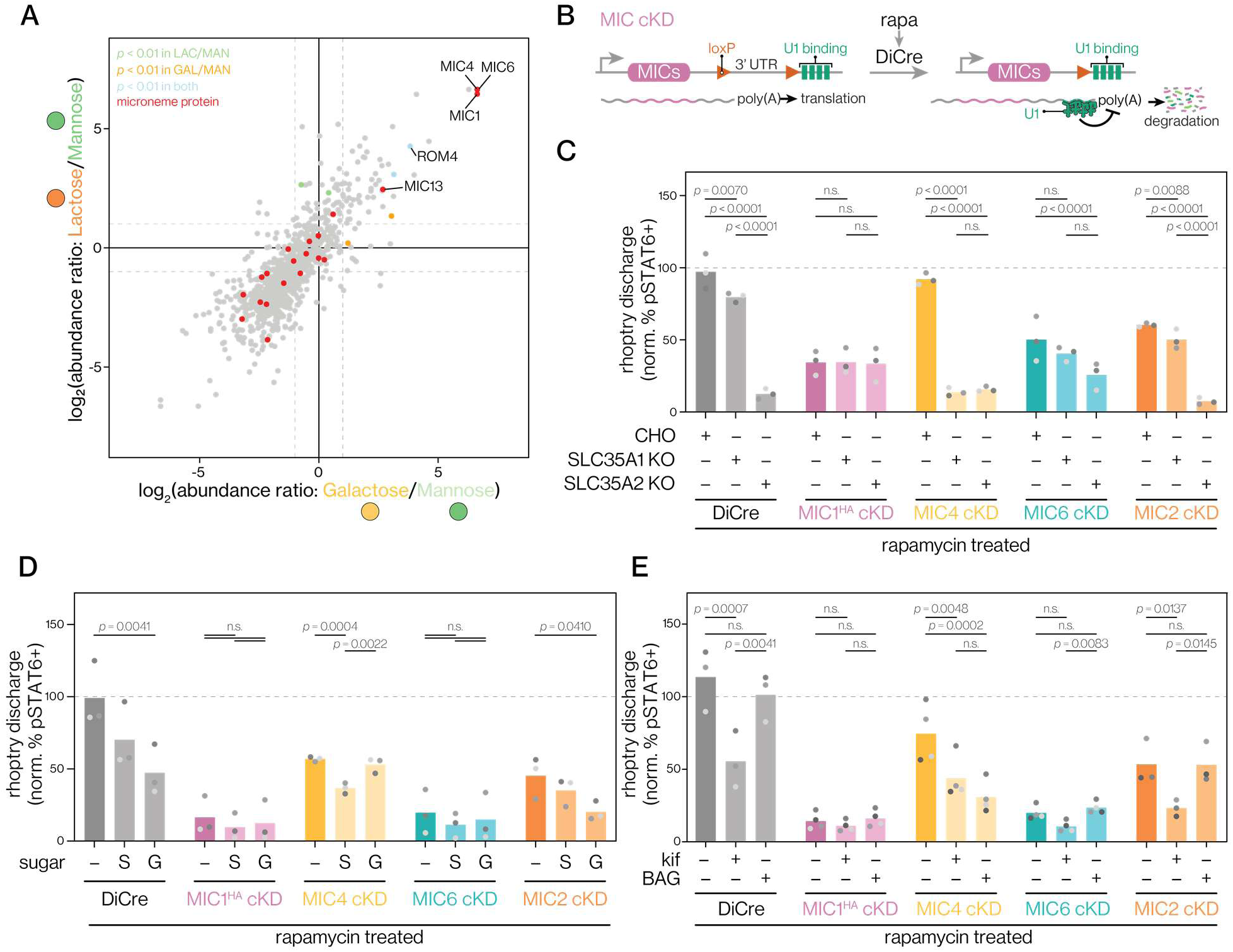
The MIC1/4/6 complex drives interaction with galactose- and sialic acid– containing N-glycans during rhoptry discharge. **(A)** Proteins from *T. gondii* lysates were differentially enriched using galactose-, lactose-, or mannose-coated agarose beads and their abundance and identity was determined by mass spectrometry. The mannose bead condition was used as the negative control for background subtraction. Dashed lines indicate a relative 2-fold enrichment. Mean of *n* = 2 biological replicates is plotted; *p*-values are derived from Benjamini-Hochberg adjusted *t*-tests and onl displayed for proteins with enrichment ratios greater than 2 in the indicated dimension. Known microneme proteins are highlighted in red. **(B)** Conditional DiCre/U1 knockdown approach used for MIC cKD strain construction replaces the endogenous 3′ UTR with a floxed 3′ UTR ahead of a tandem array of U1 motifs. Rapamycin treatment dimerizes DiCre, leading to 3′ UTR excision, replacement with the U1 array, and U1-mediated mRNA transcript degradation. **(C)** The interaction of the MIC1/4/6 complex with host galactose (SLC35A1 KO) and sialic acid (SLC35A1 KO) transport knockouts was evaluated by incubating CHO KO cell lines with rapamycin-treated DiCre parental strain or cKD strains for MIC1^HA^, MIC4, MIC6, and MIC2. Rhoptry discharge was assessed by the relative percent of pSTAT6–positive nuclei. Mean of *n* = 3 biological replicates is plotted; *p*-values derived from analysis of each parasite strain separately (vehicle and knockdown conditions) using two-way ANOVA with Tukey’s post hoc multiple comparison test. **(D)** The interaction of the MIC1/4/6 complex with exogenously added sugars evaluated by incubation of human fibroblasts with rapamycin-treated DiCre parental strain or cKD strains for MIC1^HA^, MIC4, MIC6, and MIC2. 10 mM of the indicated exogenous sugar was added with the parasites (G, galactose; S, sialic acid). Rhoptry discharge was assessed by the relative percent of pSTAT6–positive nuclei. Mean of *n* = 3 biological replicates is plotted; *p*-values derived from analysis of parasite each strain separately (vehicle and knockdown conditions) using two-way ANOVA with Tukey’s post hoc multiple comparison test. **(E)** The interaction of the MIC1/4/6 complex with N- and O-glycosylation evaluated by incubating rapamycin-treated DiCre parental strain or cKD strains for MIC1^HA^, MIC4, MIC6, and MIC2 with human fibroblasts pretreated for 4 days with kif (10 μM) or BAG (2 mM). Rhoptry discharge was assessed by the relative percent of pSTAT6–positive nuclei. Mean of *n* = 3–4 biological replicates is plotted; *p*-values derived from analysis of each parasite strain separately (vehicle and knockdown conditions) using two-way ANOVA with Tukey’s post hoc multiple comparison test.

### The MIC1/4/6 complex binds host galactosyl N-glycans required for efficient rhoptry discharge

Prior work found that MIC1 and MIC4 bind sialic acid and galactose, respectively ^76,77,84^. The transmembrane domain–containing MIC6 anchors the complex to the parasite plasma membrane but does not contain glycan-binding domains ^83^. The MIC1/4/6 complex has been shown to contribute to invasion but not attachment, a phenotype consistent with a role in rhoptry discharge ^85^. To directly examine the role of these microneme proteins in rhoptry discharge, we generated conditional knockdown (cKD) strains for all members of the complex, as well as for the control microneme protein MIC2, which contributes to parasite attachment and binds heparin ^22,35,36^. We used the U1 transcriptional cKD system, where the endogenous 3′ UTR of a target gene is replaced with a floxed 3′ UTR followed by four tandemly arrayed U1-binding sites ^44^. When generated in the DiCre background ^43^, a 2 h rapamycin treatment leads to excision of the 3′ UTR, blocking proper mRNA processing and expression of the target protein (**Figure 4B**). As part of constructing the MIC1 cKD, we included a C-terminal HA tag, which demonstrated robust knockdown of the microneme-localizated HA signal upon rapamycin treatment (**Figure S4A**). Epitope tagging was avoided for MIC4 and MIC6 because we suspected it would interfere with their function. However, knockdown of MIC2 could also be confirmed with a specific antibody (**Figure S4B**).

We then examined the genetic interactions between the disrupted host and parasite pathways, looking for non-additive effects (**Figure 1D**). In the absence of rapamycin treatment, all cKD strains showed identical rhoptry discharge profiles across all the CHO cell lines tested (**Figure S4C**). However, upon knockdown, we observed clear strain-specific effects. MIC2 knockdown showed an additive effect with the loss of host sialylation (SLC35A1 KO) or galactosylation (SLC35A2 KO), indicating that host cell adhesion sets the overall levels of rhoptry discharge but does not interact directly with these N-glycosylation pathways (**Figure 4C**). By contrast, knockdown of MIC1 clearly showed non-additive effects with the host mutations, indicating that it is crucial for the effect of host N-glycans on rhoptry discharge. Knockdown of MIC6 had a similar effect, displaying only subtle additivity. Surprisingly, despite MIC4’s role in galactose binding, knockdown had a minimal effect on rhoptry discharge into wild-type CHO cells. However, knockdown strongly compounded with the loss of host sialylation (SLC35A1 KO). These results agree with the organization of the MIC1/4/6 complex and the reported glycan specificities of MIC1 and MIC4. For example, MIC1 mediates attachment of MIC4 to the complex, so loss of MIC1 or MIC6 should render parasites unable to bind either sialic acids (via MIC1) or galactose (via MIC4). By contrast, MIC1 and MIC6 still associate when MIC4 is lost ^76,83,84,86^), so the complex can lose galactose binding while retaining sialic-acid binding— consistent with the extreme reliance of the MIC4 mutants on host sialylation (SLC35A1 KO), which is also lost when galactosylation is impaired (SLC35A2 KO).

We also examined the role of the MIC1/4/6 complex on rhoptry discharge into human fibroblasts, measuring the effect of adding exogenous sugars on this process. All of the strains displayed similar levels of rhoptry discharge without rapamycin treatment, which were reduced by exogenous sialic acid and, to a greater degree, galactose (**Figure S4D**). As with the CHO mutants, MIC2 knockdown reduced rhoptry discharge, but was further inhibited by the exogenous sugars (**Figure 4D**). By contrast, loss of MIC1 or MIC6 strongly inhibited rhoptry discharge into human fibroblasts, with no additional effect from the addition of sialic acid or galactose. Consistent with the specific loss of galactose recognition by the MIC1/4/6 complex, MIC4-deficient parasites were modestly affected by exogenous sialic acid but not galactose.

Recombinant MIC1 and MIC4 have been shown to bind N-glycans ^76,77,82^. To confirm whether the MIC1/4/6 complex indeed regulates rhoptry discharge through N-glycan binding, we used kif to reduce the levels of host cell complex N-glycans, relying on the reduction of host O-glycans by BAG as a control. As expected, kif treatment had no effect on the already suppressed rhoptry discharge of the MIC1 and MIC6 knockdowns, but had additive effects with knockdown of MIC2 or MIC4 (**Figure 4E, S4E**). These results are consistent with the complete loss of N-glycan binding by the complex when either MIC1 or MIC6 are lost, but the partial retention of sialic acid binding by the MIC1/6 sub-complex when MIC4 is disrupted. Intriguingly, BAG treatment only inhibited rhoptry discharge when MIC4 had been disrupted, suggesting that either the MIC1/6 subcomplex may secondarily bind O-glycans or, more likely, that NeuAcα2-3 linkages—the dominant glycan thought to be bound by MIC1—are suppressed by the known off-target activity of BAG ^87,88^. Taken together, these results point to MIC1/4/6 complex as the major mediator of host N-glycan recognition during rhoptry discharge, specifically through the redundant MIC1 recognition of host sialylation and MIC4 recognition of galactosylation.

### The MIC1/4/6 complex drives cholesterol-dependent formation of membrane microdomains

We considered the relationship between the recognition of host N-glycosylation by the MIC1/4/6 complex and host cholesterol, the second major pathway identified in our screen for rhoptry discharge factors (**Figure 1C**). By analogy, clathrin-independent carrier (CLIC) endocytosis (also known as GEEC) relies on human galectin-mediated clustering of N-glycosylated proteins into microdomains, which depends on the presence of cholesterol in the plasma membrane ^89,90^. Intriguingly, CLIC/GEEC has been shown to play prominent roles in infection and immunity ^91,92^. We hypothesized that the MIC1/4/6 complex acts analogously to galectins, clustering N-glycosylated proteins in a cholesterol-dependent manner to mediate efficient rhoptry discharge. To test this hypothesis we measured rhoptry discharge in the various MIC knockdowns under varying levels of host plasma membrane cholesterol. We depleted host cell membrane cholesterol as we had previously done by MβCD extraction followed by varying levels of reconstitution and increased host cell membrane cholesterol by loading host cells with excess cholesterol (**Figure 5A**). Following membrane cholesterol modulation, host cells were challenged with parasites that had been treated with either vehicle or rapamycin to induce knockdown of the various MIC proteins (**Figure 5B–F, S5A-B**). As anticipated from our earlier experiments, lowering host cholesterol decreased rhoptry discharge by the parental DiCre strain, while increasing host cholesterol had a negligible effect and rapamycin did not affect these trends (**Figure 5B**). Similar trends were observed when knocking down MIC2 or MIC4 (**Figure 5D and F, S5A–B**). By contrast, knockdown of MIC1 or MIC6 sensitized parasites to lowered cholesterol levels, whereas increasing host cholesterol significantly enhanced rhoptry discharge (**Figure 5C and E, S5A-B**). Together, these results suggest the MIC1/4/6 complex is orchestrating clustering of N-glycosylated proteins and cholesterol to mediate host recognition or pore formation for rhoptry discharge (**Figure 5G**).

**Figure 5.**
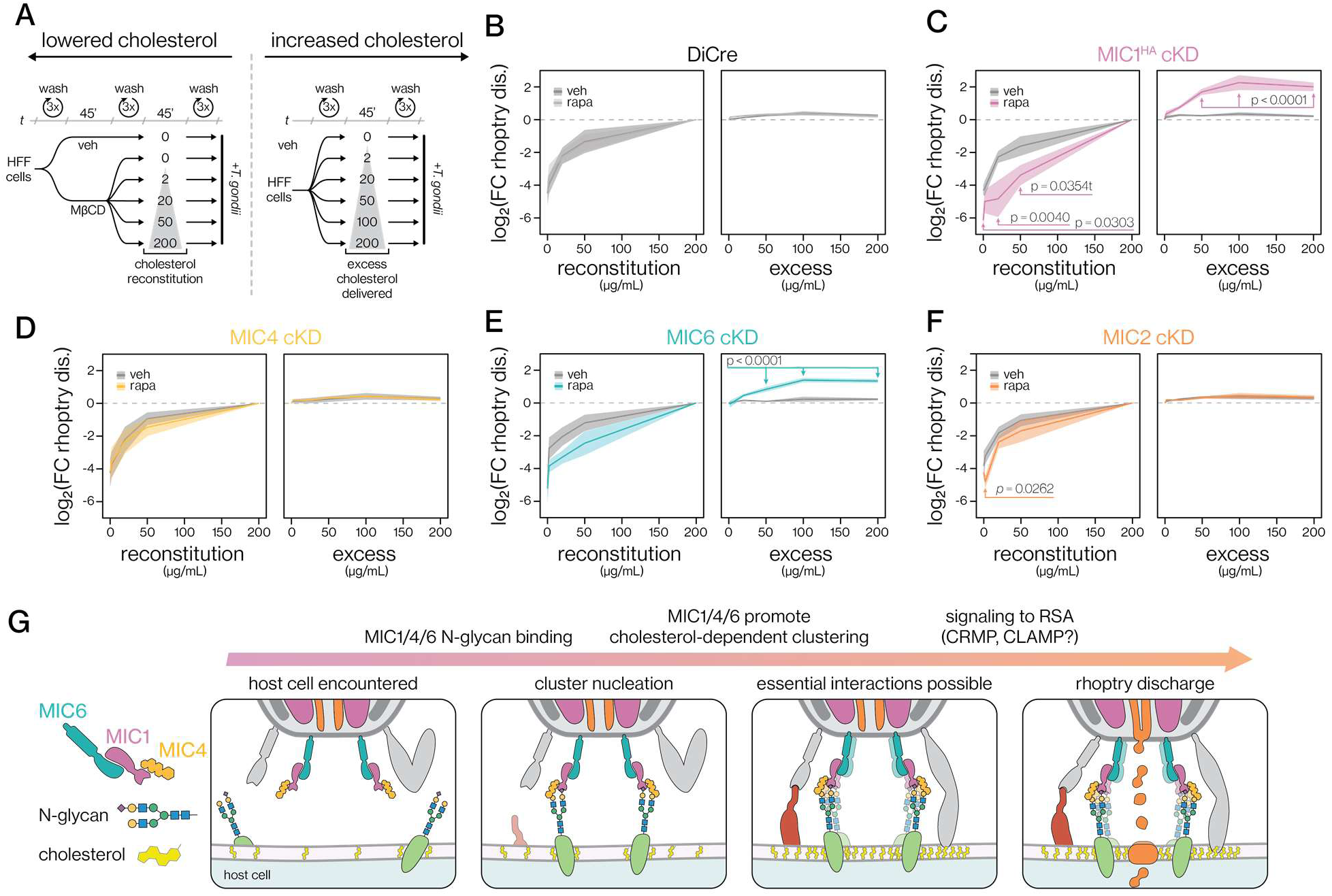
The activity of the MIC1/4/6 complex is sensitive to host cholesterol levels in rhoptry discharge. **(A)** To modulate cholesterol levels, human fibroblasts were depleted of cholesterol by a 45-min treatment with MβCD. Varying concentrations of water-soluble cholesterol were then used to reconstitute cholesterol into host cells for 45 min. To increase host cholesterol, untreated human fibroblasts were loaded with indicated concentrations of extra water-soluble cholesterol for 45 min. Following the various treatments, host cells were challenged with parasites. **(B-F)** Rhoptry discharge into cholesterol-depleted or supplemented condition was evaluated following challenge with DiCre parental **(B)** or cKD strains for MIC1^HA^ **(C)**, MIC4 **(D)**, MIC6 **(E)**, or MIC2 **(F)**, pre-treated with vehicle or rapamycin to induce recombination. Rhoptry discharge was assessed by the relative percent of pSTAT6–positive nuclei, then normalized to the highest cholesterol reconstitution level (left panels) or the vehicle-treated condition (right panels) prior to calculation of relative fold-change and log_2_ transformation. Mean of *n* = 3 biological replicates is plotted with error bars indicating s.e.m. *p*-values derived from analysis of each parasite strain separately using two-way ANOVA with Tukey’s post hoc multiple comparison test. **(G)** Model for the role of host N-glycans, cholesterol, and their coordination of rhoptry discharge into host cells by the MIC1/4/6 microneme complex. Parasite factors involved in other host cell interactions are displayed in grey.

## DISCUSSION

As apicomplexan proteins required for rhoptry discharge have been discovered, the role of the host cell in the process has remained elusive. We performed an unbiased host-directed screen that narrowly focused on the essential step of rhoptry discharge. The screen uncovered host galactose-containing N-glycans and cholesterol as specifically required for *T. gondii* rhoptry discharge. Analyzing dual perturbations in the parasite and host, we conclude that these pathways act parallel to known rhoptry discharge factors, including the CLAMP and CRMP complexes ^10–12,36^. Instead, we find that the previously discovered MIC1/4/6 microneme complex plays a novel role in rhoptry discharge by binding galactose-containing N-glycans on the host plasma membrane. Based on the observed cholesterol dependency of these interactions, we propose that the MIC1/4/6 complex nucleates the formation of microdomains rich in N-glycans and cholesterol. The induced microdomains are likely required for downstream steps in rhoptry discharge, potentially promoting receptor binding by the CLAMP and CRMP complexes or establishing the mechanical requirements for pore formation during rhoptry-discharge induced pore formation. Our work represents a novel framework through which to understand conserved and divergent aspects of apicomplexan rhoptry discharge.

Rather than a specific protein providing the crucial receptor, our screen found that rhoptry discharge depends on N-glycosylation. Two of the strongest effects observed in our screen were caused by disruption of *UGP-2* and *SLC35A2* which are required for the biosynthesis and golgi transport of UGP-galactose for galactosylation of lipids and proteins. Galactosylation could be shown to be crucial in rhoptry discharge as knockout of SLC35A2 or UGP-2, or the exogenous addition of galactose-containing sugars, severely decreased rhoptry discharge. The role of galactosylation could be specifically localized to complex-type N-glycans based on the effect of treatment with kifunensine—a specific inhibitor of complex N-glycan maturation ^65,66^— which excluded other galactose containing glycans like GSLs and O-glycans ^52^. Complex N-glycans are typically capped with either galactose or sialic acid, yet knocking out the sialic acid transporter SLC35A1 only modestly affected rhoptry discharge. The dominant effect of galactosylation can be attributed to its requirement for sialic acid addition to N-glycans. As the MIC1/4/6 complex can redundantly bind both galactose and sialic acid, both interactions must be inhibited to completely disrupt the function of the complex. This dual inhibition is achieved by either genetic removal of galactosylation (e.g. SLC35A2 knockout) or combined disruption of sialylation (SLC35A1 knockout) with exogenous galactose treatment. Furthermore, when the galactose-binding MIC4 is knocked down, this redundancy is broken and rhoptry discharge becomes highly dependent on sialylation. Redundant binding of galactose and sialic acid by the MIC1/4/6 complex might contribute to the broad tissue tropism of *T. gondii*, by accounting for the differences in glycosylation throughout host cell types.

Cell surface glycans are hot spots for host-pathogen interactions ^93^, and different glycans can dramatically influence susceptibility and tropism. For influenza, the abundance of α2,6 and α2,3-linked sialic acids in airway epithelial cells represents a key determinant of host species tropism, impacting the differential susceptibility of humans and birds to different viruses ^94,95^. Host glycans also mediate bacterial attachment and infection, such as oligomannose N-glycan recognition by the FimH adhesin of uropathogenic *E. coli* ^96^. Pathogen glycans can conversely drive attachment, as occurs with *Chlamydia trachomatis* glycans that are bound by host galectin-1 to drive attachment and infection in the genital tract ^97^.

The glycans bound by *T. gondii* through the MIC1/4/6 complex are broadly distributed across mammalian cell types. MIC1 and MIC4 bind N-glycans with α2,3-linked sialic acid and terminal galactose, respectively. Such glycans are found on cells in every tissue, though their relative abundances vary between cell types and species ^76,77,82,98,99^ the differences between CHO cells and human fibroblasts in our work. For example, MIC4 disruption has a negligible impact on rhoptry discharge into CHO cells but reduces rhoptry discharge into human fibroblasts by ∼50%. The difference between cell types may be due to the overall density of glycans—a hypothesis supported by the relatively more pronounced effect of exogenous glycan competitions on rhoptry discharge into human fibroblasts. Interestingly, the malarial parasite *Plasmodium falciparum* uses the *Pf*RCR complex to bind α2,6-linked sialic acid N-glycans, which might dictate the narrower tropism for erythrocytes ^41,100^. *T. gondii*’s preference for α2,3-linked sialic acids may contribute to its broader tropism and enable different modes of interaction with host cells. Mammalian Galectin-3 binds 2,3-linked sialylated N-glycans to form microdomains—as we have proposed for MIC1/4/6—and its binding is blocked by α2,6-linked sialylation ^102–104^.

The MIC1/4/6 complex was previously implicated in glycan binding ^76,77,81,82^, and this is further corroborated by our work. Based on affinity to lactose- and galactose-coated beads, we identified only the strongest interactors—capable of withstanding stringent washing—which prominently included the MIC1/4/6 complex. Previous studies characterized the sugar-binding profile and assembly of the MIC1/4/6 complex using recombinant and parasite-derived proteins ^76,77,81,82^. The MAR domain of MIC1 has a clear preference for α2,3-linked sialic acids ^76^. MIC4 has six Apple domains with the fifth being responsible for binding both β1,3- and β1,4-linked galactose ^77^. MIC6 lacks lectin domains, but has a transmembrane helix that helps anchor the rest of the complex to the parasite plasma membrane. Via its EGF domains, MIC6 binds MIC1’s galectin-like domain, while MIC1, in turn, tethers MIC4 by binding its first two Apple domains ^83,105^. We therefore conclude that binding of the complex to galactose and lactose beads occurs through MIC4. Tight galactose binding by MIC1/4/6 might be related to the proposed higher-order assemblies of the complex ^77^. Other parasite lectins may remain to be discovered, since we have not profiled sialic acid–binding or all possible galactose linkages and conformations. However, our genetic interaction studies indicate that MIC1 and MIC4 redundantly control the major interactions with host N-glycans during rhoptry discharge. The general binding of N-glycans by MIC1 and MIC4 may play additional roles during infection, which would act parallel to the functions in rhoptry discharge presented here. In certain cell types, N-glycan binding by MIC1 and MIC4 activates innate immunity signaling via Toll-like receptor (TLR) activation ^82^. This effect can be induced by recombinant MIC1 and MIC4 and disruption of MIC1 limits the induction of pro-inflammatory cytokines in murine infection models, leading to reduced cytokine production and extended host survival. It is unclear whether this response is a specific adaptation to targeting TLRs or related to the effect of N-glycan clustering on specific cell types. Indeed, clustering can be a general mechanism for activation across a wide array of receptors ^106^ induction of pro-inflammatory signaling may drive immune pressure, which has been associated with chronic differentiation of parasites ^107,108^. Different attachment and host recognition strategies throughout tissues may affect global host responses in ways that alter the course of infection.

In addition to N-glycans, our screen identified cholesterol biosynthesis as a second major host factor involved in rhoptry discharge. Cholesterol is an essential plasma membrane lipid that regulates membrane fluidity and participates in various processes that cluster membrane components or reshape the membrane ^57^. Cholesterol is also required for *T. gondii* invasion ^38,39^, appearing to be enriched at the invasion site, then lost from the parasitophorous vacuole ^38,54^. Early work also implicated host cholesterol in rhoptry discharge by following the translocation of certain markers by microscopy ^38,39^. Removal of cholesterol from erythrocytes analogously blocks *P. falciparum* invasion, suggesting that host cholesterol contributes to invasion of other apicomplexans ^109,110^. In our study, we modulated host cholesterol levels using MβCD, a cyclodextrin commonly used to extract cholesterol ^38,39,111^. Due to its essential functions in membranes, directly manipulating cholesterol can damage cells. We excluded such effects by using short treatments, consistent with previous protocols, and, crucially showing that reconstitution of cholesterol specifically and completely rescued rhoptry discharge. Our results demonstrate that rhoptry discharge depends on the abundance of host membrane cholesterol. Depletion of host cholesterol sensitizes parasites to loss of the MIC1/4/6 complex, whereas excess cholesterol alleviates the loss of MIC1 and MIC6, which suggests that N-glycan binding and cholesterol orchestrate related events in rhoptry discharge.

Cholesterol may play several roles in rhoptry discharge. We propose that MIC1/4/6 participates in microdomain nucleation, analogously to clathrin-independent endocytosis (CLIC/GEEC) in mammalian cells. CLIC/GEEC depends on galactose-containing N-glycan binding by oligomeric galectin-3 to cluster glycoproteins into cholesterol-rich microdomains, which induces membrane bending and bulk endocytosis ^89,91^. Galectins in CLIC/GEEC can also be inhibited by exogenous lactose or cholesterol depletion ^89,112,113^. Alternatively, host cholesterol may need to be concentrated to signal contact with host cells through other parasite proteins (e.g. CLAMP, CRMP, or MIC8 complexes), either directly or indirectly through the accumulation of other lipids or proteins. Conversely, cholesterol may simply facilitate protein mobility to or stability within microdomains. Supporting such microdomain formation, proteins enriched in cholesterol-dependent lipid rafts are generally incorporated into the parasitophorous vacuole, along with cholesterol and the sphingolipid Gm1 ^38,105,114,115^. Cholesterol also alters the mechanical properties of membranes, which may be required for membrane deformation or translocation of rhoptry proteins, but might not affect signaling of rhoptry discharge per se. It should be noted that rhoptry discharge into the extracellular space has not been observed, although this may be attributed to the sensitivity of current experimental methods. An intriguing parallel may be the cholesterol-dependent cytolysin toxins’ use of glycan recognition in combination with cholesterol to form pores in the target plasma membrane ^116^; rhoptry-driven perforation may have similar biophysical constraints. A more precise understanding of cholesterol’s role in rhoptry discharge will require the development of biophysical methods to distinguish between these possible roles.

Another open question is the interplay between clade-specific factors like MIC8 and the MIC1/4/6 complex, and the pan-apicomplexan CRMP and CLAMP complexes. The clade-specific factors might reflect adaptation to specific niches, either at the level of individual species or for different stages in their life cycles. The MIC1/4/6 complex is down-regulated in chronic bradyzoite stages of *T. gondii* ^117,118^, which are thought to be adapted to transmission ^119^ and display different cell type tropism ^120^. Differential reliance on MIC1/4/6 between acute-stage tachyzoites and chronic-stage bradyzoites would support the notion that this complex facilitates invasion of a wider array of host cells by tachyzoites. *P. falciparum* depends on clade-specific N-glycan–binding proteins, such as the aforementioned *Pf*RCR, to invade erythrocytes ^41^. Under the proposed model, the microdomains formed by the clade-specific parasite factors may be required for the function of pan-apicomplexan factors or may simply act as parallel requirements for rhoptry discharge. This model is consistent with the additive effects we have documented for the perturbation of host N-glycosylation or cholesterol and genetic disruptions affecting the CLAMP or CRMP complexes.

In summary, we discovered that the MIC1/4/6 microneme complex coordinates host N-glycans and cholesterol to form a microdomain rich in lipids and proteins. We propose that these microdomains support efficient host cell interactions by other microneme complexes or provide a mechanical scaffold that facilitates rhoptry discharge and parasite invasion. The key role of these interactions in parasite invasion suggests they could be therapeutic targets. The MIC1/4/6 complex has been tested as a potential vaccine target. Immunization with purified MIC1 and MIC4 produced a protective response against *T. gondii* infection^121^. Additionally, a strain lacking both MIC1 and another microneme protein, MIC3, protected mice against a subsequent challenge, suggesting it could be used as a live-attenuated vaccine^122^. Targeting parasite-host glycan interactions with small-molecule inhibitors has recently shown promise in experiments with *P. falciparum* ^41^. Beyond its therapeutic potential, this work refines our mechanistic understanding of rhoptry discharge, providing a conceptual framework to evaluate the hierarchy of the other host-parasite interactions required for invasion.

## MATERIALS & METHODS

### Mammalian cell culture and genetic manipulations

Human foreskin fibroblast (HFF) cells were expanded in “D10”: DMEM (Gibco, 11965118) with 10% serum (heat-inactivated foetal serum [IFS]; Sigma, F4135-500ML). For maintenance of confluent flasks and *T. gondii* passaging, DMEM with 3% newborn calf serum (Sigma, N4762-500ML) was used in place of fetal serum (D3C). All complete DMEM media was supplemented with 2 mM L-glutamine (Gibco, A291680) and 10 μg/mL Gentamicin (Gibco, 15710072). K562 cells (Cellosaurus: CVCL_0004) were grown in “R10”: RPMI-1640 medium (Gibco, 11875085) supplemented with 10% IFS (Sigma, F4135-500ML), 2 mM L-glutamine, and 10 μg/mL Gentamicin. CHO cells – Pro-5 (parental; Cellosaurus: CVCL_4382), Lec1 (MGAT KO), Lec2 (SLC35A1 KO), Lec8 (SLC35A2 KO) – were all grown in Ham’s F-12 Nutrient Mix (Gibco, 11765047), supplemented with 10% IFS, 2 mM glutamine, and 1:100 Penicillin-Streptomycin (Gibco, 15140-122). All cells were maintained at 37**°**C and 5% CO_2_.

To generate UGP-2 KO and SLC35A2 KO K562 cells, guides targeting the coding were cloned into the pX458 vector expressing *Sp*Cas9-T2A-eGFP (Addgene Plasmid #48138) to generate plasmids pDV046_UGP2_3_sgRNA_pX458 and pDV045_SLC35A2_3_sgRNA_pX458. 20 µg of these plasmids was heat-sterilized (65°C, 15 min) and combined with 4×10^6^ K562 cells in 500 μL of serum-free RPMI-1640 and transferred to a 4 mm electroporation cuvette (BTX, 45-0126) on ice. A ECM-830 square wave electroporator (Harvard Instruments, BTX) was used to electroporate the cells (Low Voltage Mode, 250 V, 9 msec pulse length, field strength 625V/cm). Transfected cells were immediately transferred to complete R10 medium. After 24 h recovery, GFP-expressing cells were sorted into 96 well plates and grown for 12-14 days, then growing clones were transferred to 12-well plates. The lectin GS-II from *Griffonia simplicifolia* conjugated to Alexa Fluor 647 (Invitrogen, L32451), which binds terminal GlcNAc, was used to positively screen for clones clones for successful knockouts of UGP-2 and SLC35A2 ^61,123^. Cells were washed once with FACS buffer (PBS with 1% serum, 1 mM CaCl_2_, 1 mM MgCl2, and 1 mM MnCl2), stained with 5 μg/mL GS-II in FACS buffer for 15-30 min on ice then resuspended in lectin-free FACS buffer and analyzed using flow cytometry using a MACSQuant VYB flow cytometer (Miltenyi Biotec).

K562 cells with UGP-2 KO (clone 3C.2) and SLC35A2 KO (clone 2B.1) were grown the same way as parental K562 cells. For galactose recovery experiments K562 cells were grown in R10 supplemented with 25 mM galactose for 3 days (or a water vehicle control). To assess the change in surface glycosylation patterns in presence of galactose, GS-II staining and flow cytometry was performed as above at the same time point as the rhoptry discharge assays.

### *T. gondii* genetic manipulations

To generate the DiCre/MyoH-U1/Toxofilin-βla strain, a HXGPRT-Toxofilin-βla repair template was amplified from a previously described vector used to generate transgenic Toxofilin-βla strains ^34^ using primers oEB080 and oEB081 containing homology arms for a defined neutral genomic locus ^124^. The repair template was co-transfected with a gRNA/Cas9 containing plasmid targeting the neutral locus for double stranded break ^125^ then selected for integration with 200 ug mycophenolic acid (MPA) and 50 ug xanthine (XA). After selection, clonal parasites were isolated by limiting dilution.

To generate RH/Δ*ku80*/Δ*hxgprt*/dTomato (TgDV101), primers oALH091 and oALH092 were used to amplify a dTomato with a pTub1 promoter and a DHFR 3’utr (from pBM017) with homology arms for the neutral locus. Parasites were transfected with a gRNA/Cas9 containing plasmid targeting the neutral locus, and the next day single parasites were sorted by FACS into 96 well plates. The dTomato SPATR knockout strain – RH/Δ*ku80*/Δ*hxgprt*/Δ*spatr/*dTomato – was generated with same way from a previously constructed RH/Δ*ku80*/Δ*hxgprt*/Δ*spatr* strain ^58^.

The strain background for U1 conditional knockdowns is RH/Δ*ku80*/Δ*hxgprt*/DiCre/Ty-APH, and is referred to as “DiCre” throughout. This strain was generated in the previously characterized stable DiCre background ^43^ by co-transfecting a PCR-amplified template containing the Ty tag sequence with homology arms to the N-terminus of the the microneme surface protein APH alongside a gRNA/Cas9 plasmid that with a gRNA targeting the N-terminus of the *APH* coding sequence. A clonal population was isolated via limiting dilution without prior selection. To generate conditional knockdown strains using the U1 system, we used HiT vectors ^126^ that target a construct with the 3’ UTR of CDPK3 between loxP sites followed by 4xU1 motifs to the C-terminal codon of the target gene ^44^. In this system, rapamycin addition leads to excision of the CDPK3 3’ UTR and appends the U1 motifs, such that the mRNA is targeted for mislocalization and degradation, and leading to protein knockdown. The HiT vectors used for making MIC1^HA^ cKD, MIC4 cKD, and MIC2 cKD strains contained a mNeonGreen fluorescent protein cassette as a selectable marker and were isolated by single parasite FACS sorting. The MIC6 cKD strain was made instead by PCR-amplification of a homologous recombination template containing the same U1 regulatory elements but instead using a DHFR selectable marker, so pyrimethamine selection and isolation by limiting dilution was used to isolate clonal parasites. For all strains, parasites were validated by PCR. Where possible – for MIC1^HA^ cKD and MIC2 cKD – knockdown was additionally validated by immunofluorescence assays (IFA). For IFAs, parasites were treated with 50 nM rapamycin when added to HFF cells, followed by rapamycin washout 2 hours later. 2 days later, infected HFF cells were fixed and permeabilized with 100% -20°C methanol for 8 minutes, and blocked with blocking buffer (5% IFS and 1% normal goat serum in PBS). Samples were immunostained with blocking buffer containing 1:1000 mouse anti-HA (Biolegend #901501), 1:1000 mouse anti-MIC2 ^127^, 1:10000 guinea pig anti-CDPK1 ^118^. After 3x PBS washes, samples were stained with anti-mouse Alexa Fluor 488 (A-11029, Thermo), anti-guinea pig Alexa Fluor 647 (A-21450, Thermo), and 1:20000 Hoechst 33258 (sc-394039, Santa Cruz). Samples were imaged on a Nikon CSU-W1 spinning disk confocal microscope using a 60X silicone oil objective and processed using Fiji ^128^.

To generate the CRMPa cKD strain, a HiT vector ^126^ targeting the C-terminus of CRMPa (TGGT1_261080) with a V5-TEV-mNeonGreen-mAID-Ty tag and the 3’ UTR of CDPK3 was generated. Alongside a Cas9 transient expression vector, this HiT vector was transfected into an RH/Tir1 strain background that heterologously expresses the F-box protein Tir1 such that indole-3-acetic acid (IAA) addition induces the degradation of mAID-tagged proteins ^126,129,130^. The HiT vector contained a DHFR selection cassette and pyrimethamine was used to select for its integration into the CRMPa locus, then correct integration was validated using PCR. For all experiments using this RH/Tir1/CRMPa cKD (CRMPa cKD) strain, freshly lysed out parasites were added to fresh HFF monolayers and allowed to invade for 1 hour. 500 µM IAA or a PBS vehicle control was then spiked into the flasks to knock down CRMPa as indicated, and the parasites were grown for 2 more days.

### Host cell rhoptry discharge screen

For the host cell rhoptry discharge screen, K562 cells were maintained in shaking flasks in an Infors HT incubator at 37**°**C, 125 RPM, and 8% CO_2_ with an added 0.1% pluronic F-68 (Thermo) to prevent cell clumping and increase viability during mechanical stress from shaking.

The screen was performed with 2 host cell library pools, infected 3 weeks apart, with 2 replicates of the rhoptry discharge performed on each host cell pool, resulting in 4 assay replicates. Throughout library generation and selection, host cells were maintained at a targeted minimum of 1000x library coverage.

To generate the host cell library, a pooled human genome-wide optimized 5-guide library (97,888 unique guides) packaged into lentivirus prepared by the Whitehead Genome Technology Core was used to infect K562 host cells ^45^. Viral transduction was performed by spinfecting 300 million K562 cells in 6 well dishes with 5 million K562 cells/well at an MOI of 0.5 (titered functional transduction units : host cell) at 1200 g for 45 minutes at 37°C in the presence of 8 μg/ml polybrene. The next day, the media was refreshed. 2 days after lentivirus infection, cells were resuspended in medium containing 2 μg/mL puromycin (Sigma) to select for guide vector incorporation. 5 days after lentiviral infection, puromycin was removed, and cells were counted to ensure library coverage was maintained. On day 13 and 14, 350-450 million host cells were used to perform sorting in the rhoptry discharge assay, and 150 million untreated host cells divided into aliquots of 50 million cells were harvested to sequence the library diversity at the time of each assay replicate.

To generate the parasites for each replicate of the rhoptry discharge assay, 2 days prior (i.e. day 11 and 12), 70 15-cm dishes of HFF cells were each infected with 3.5e7 DiCre/MyoH-U1/Toxofilin-βla parasites and treated with 50 nM rapamycin in D3C medium. A minimum of 2 hours later, rapamycin was washed out by 2 washes of PBS, incubation in warmed D3C for >5 min, followed by a third PBS wash. Fresh warm D3C was added to each plate, and the plates were returned to the incubator for approximately 42 hours. On the day of each rhoptry assay replicate, the plates were scraped and parasites were released from host cells by syringe-release using 27 gauge needles. The parasites were filtered through 5 μm pore filters to remove debris and unlysed cells, and pelleted by centrifugation at 1,000 g for 5 minutes. All tubes of parasites were pooled and washed in HBSS/HEPES (HBSS with 25 mM HEPES pH7.4), pelleted again, and resuspended to a concentration of 400 million parasites/mL in HBSS/HEPES.

For each replicate of the rhoptry discharge assay, 350-450 million K562 cells were pelleted in a 500 mL conical tube, washed with HBSS/HEPES to remove serum and resuspended to 20 million cells/mL. K562 cells were mixed 1:1 with DiCre/MyoH-U1/Toxofilin-βla parasites (400 million/mL) to yield a multiplicity of parasites:host cells of 20:1 in a 50 mL conical tube and placed in a 37°C water bath. Approximately 8 mL of 6x solution of the CCF4-AM substrate was made up according to the kit guidelines with minor modifications (Thermo cat. #K1095). Briefly, 14.4 μL of solution A (1 mM CCF4-AM) was mixed with 480 μL of solution B (100 mg/mL Pluronic^®^-F127 surfactant in DMSO with 0.1% acetic acid), then 7.51 mL of HBSS/HEPES was added. After 1 h of incubation, the host:parasite mixture was allowed to cool to room temperature, then the 6x CCF4-AM loading solution was added to yield a 1X CCF4-AM concentration. After a 30 minute incubation at room temperature with gentle mixing, the cells were strained through a 50 mL conical cell strainer and placed on ice.

The host cells were sorted based on cleavage of the FRET CCF4-AM substrate into a rhoptry discharge positive (“blue”) and rhoptry discharge negative (“green”) population using two BD FACSAria IIIu cell sorters equipped with a 405 laser line and 450/40 filter. The cells were sorted using 4-way purity into 15 cm conicals pre-coated with BSA (coated by overnight incubation with 5% BSA in PBS). We targeted sorting a minimum of 1000x library coverage, or 100 million host cells in each replicate (R1: 90 mil, R2: 140 mil, R3: 140 mil, R4: 165 mil). After sorting, aliquots of 50 million host cells were made, pelleted, and frozen for Illumina workup, quality control, and sequencing alongside the unsorted controls by the Whitehead Institute Functional Genomics Platform and Genome Technology Core on a Illumina NovaSeq.

### Screen analysis

We obtained approximately 320 million reads across 12 sequenced conditions (4 each of Unsorted, rhoptry discharge susceptible [blue], and rhoptry discharge resistant[green]). Data was analyzed using the MAGeCKFlute pipeline ^131^. Briefly, the pipeline was used to map sgRNAs, make count tables, minimize batch effects (using the BatchRemove script), then the MAGeCK MLE algorithm was used to identify the importance of each knockout to rhoptry discharge. For guides that targeted multiple genes, the counts for those specific guides were duplicated amongst the co-targeted genes prior to batch correction and the MAGeCK MLE analysis. The MAGeCK MLE algorithm compared each sorted population to the unsorted controls and inferred a “beta-score”, where a positive beta-score represents enrichment in the target population. To make ranked gene plots and perform GSEA we combined the scores from the resistant and susceptible populations into a single score, the “combined rhoptry score” ^131^. GSEA enrichment analysis was performed with Rstudio using the fgsea “Fast gene set enrichment analysis” package ^132^, using the reactome database pathways ^49^.

### Validation of CHO cell surface glycosylation patterns

To validate the CHO Pro-5 (parental), Lec1 (MGAT1 KO), Lec2 (SLC35A1 KO), and Lec8 (SLC35A2 KO) had the expected glycosylation patterns, CHO cells were trypsinized, washed once with FACS buffer, and stained for 15-30 min on ice with 5 μg/mL GS-II-Alexa Fluor 647 (GS-II-AF647), Wheat Germ Agglutinin-Alexa Fluor 555 (WGA-AF555; Thermo, W32464), or Concanvalin A-Alexa Fluor 488 (ConA-AF488; Thermo, C11252). After staining, cells were resuspended in lectin-free FACS buffer and analyzed using flow cytometry using a MACSQuant VYB flow cytometer (Miltenyi Biotec).

### Growth and knockdown of *T. gondii* strains

All *T. gondii* parasites were maintained by passage in HFF cells. Where noted in experiments, “WT” parasites refer to the RH/Δ*hxgprt/*Δ*ku80* background ^58^. For rhoptry discharge assays with auxin-inducible knockdowns (CRMPa cKD), 2 days prior to the experiment freshly lysed out parasites were passaged with 2-3 drops into fresh T12.5 cm^2^ flasks and allowed to invade for at least 1 hour. The media was then exchanged for D10 media containing 500 μM indole-3-acetic acid (IAA) or a vehicle control (PBS). Parasites were then harvested 40-44 hours later. For assays using the U1-based transcriptional knockdown system, 2 days prior to the experiment freshly lysed out parasites were passaged with 2-3 drops into fresh T12.5 cm^2^ flasks containing either 50 nM rapamycin to induce DiCre dimerization or a vehicle control (DMSO). After 2 hours, the flasks were washed 3X with PBS to remove rapamycin. Fresh D10 was added and parasites were harvested 40-44 hours later.

### Toxofilin-**β**la rhoptry discharge assays

Toxofilin-βla secretion assays were done similarly to the screen, with minor modifications. DiCre/MyoH-U1/Toxofilin-βla parasites were prepared as in the screen: treated with rapamycin for 2 h in T12.5 flasks, and harvested 40-44 hours later by syringe lysis using a 27G needle, followed by filtering through 5 μm pore filters to remove host debris. Parasites were isolated by centrifugation at 2000 g, resuspended in HBSS/HEPES, and counted. Parasites were then centrifuged again, and resuspended to a concentration of 1e7 parasites per 50 μL in HBSS/HEPES. To prepare host cells, K562 cells were maintained in T175 flasks such that they would be at a density of 0.5-2e6/mL at the time of assay. Their density was assessed using a Countess II Automated Cell Counter (Thermo Fisher), then they were pelleted (300-500 g, 5 min) and resuspended to a density of 5e5 K562 cells per 100 μL in HBSS/HEPES. In the wells of a 96 well tissue culture plate, 100 μL of K562 cells were then mixed with 50 μL of parasites (multiplicity of infection = 20), centrifuged for 5 min at 290 g, and incubated at 37°C for 60 min. As in the screen, 6X solution of CCF4-AM was prepared, but with propidium iodide added as a live/dead counterstain (Sigma, P4170; final concentration of 1 μg/mL). After letting the cells cool to room temperature, 30 μL of 6X CCF4-AM solution was added to each well. Samples were repeatedly pipetted to ensure cells were well mixed with CCF4-AM then incubated in the dark for at least 30 min. Each condition was assessed using a MACSQuant® VYB flow cytometer (Miltenyi Biotec). As with the screen, the level of rhoptry discharge into host cells was assessed by monitoring the ratio of host cells in the blue fluorescence channel using FlowJo (BD). Only cells that were resistant to propidium iodide staining were used in analysis. GraphPad Prism was used to perform statistical testing.

### STAT6 phosphorylation rhoptry discharge assays

All STAT6 phosphorylation (pSTAT6) rhoptry discharge assays were performed in black-walled 96 well plates with clear bottoms for imaging. HFF host cells were typically prepared by splitting a 12.5 cm confluent flask per 96 well plate and grown for 3 days prior to the experiment such that they were freshly confluent at time of the assay. CHO cells were prepared by seeding either 10,000 cells per well 3 days prior or 25,000 cells per well 2 days prior to the experiment, such that they were >90% confluent at the time of assay.

*T. gondii* parasites grown in HFF cells in T12.5 cm^2^ flasks were typically harvested immediately prior to natural egress by scraping and passing through a 27G syringe, followed by passing through disposable 5 μm pore size syringe filters to remove host cell debris. The parasites were pelleted (2000 g, 5 min), resuspended in 800 μL DMEM with 20 mM HEPES pH 7.4 added (DMEM/HEPES), transferred to 1.5 mL microcentrifuge tubes, and pelleted again. Parasites were resuspended DMEM/HEPES with 1% serum and 1.2 μM mycalolide B (Fujifilm Wako Pure Chemical Corporation; 132-12081 or Enzo; BML-T123-0020) in a volume of 100 μL per T12.5 cm^2^ flash of parasites prepared, then incubated at RT for 30-60 minutes. During this incubation, parasite density was assessed. Parasites were pelleted again and resuspended in 1 mL of D10 with 20 mM HEPES pH7.4 (D10H) for HFF experiments or Ham’s F12 with 10% serum and 20 mM HEPES pH7.4 (F12H) for CHO cell experiments. Parasites were diluted to either 1e6 parasites/mL; in the case of sugar or drug treatments during the assay, 2x working stocks 2e6 parasites/mL were prepared. Unless otherwise noted, 2e5 parasites/ well of a 96 well plate were used for all experiments.

For sugar treatments, 0.5 M stock solutions of L-galactose, D-galactose (Gal), lactose (Lac), mannose (Man), N-acetyl-galactosamine (GalNAc), N-acetyl-galactosamine (GlcNAc), and N-Acetylneuraminic Acid (Sia), were prepared in PBS. 2X working solutions were prepared in the appropriate complete medium for either HFF cells (D10 + 20 mM HEPES pH7.4; D10H) or CHO cells (Ham’s F12 + 10% serum + 20 mM HEPES pH7.4; F12H). Immediately before adding 100 μL of 2x working stocks of parasites, the growing medium was replaced with 100 μL of the 2x sugar-containing medium at room temperature. Parasites were then added and mixed with the sugar solutions by repeated pipetting, the plate was centrifuged to make parasites contact host cells (290 g, 5 min), and the plate was incubated at 37°C for 2 hours. Unless otherwise noted, final 1X concentration in all sugar experiments was 10 mM.

For cholesterol depletion and add-back experiments, recently confluent HFF cells were washed 2x with PBS and 1x with DMEM lacking any additives (D0). The media was then replaced with D0 containing 20 mM HEPES pH 7.4 (D0H) and 10 mM methyl-β-cyclodextrin (MβCD), which was prepared by directly solubilizing MβCD powder (Sigma; C4555) in D0H. The cells were incubated at 37°C for at least 45 minutes. For cholesterol add-back experiments, water-soluble cholesterol (Sigma; C4951) was dissolved in water at a stock concentration of 10 mg/mL, then the indicated concentration or vehicle control was added to D0H. After MβCD treatment, the HFF cells were washed as before, the indicated concentration of cholesterol was added and the cells were incubated again at 37°C for at least 45 minutes. The cholesterol treatments were washed out with 2x PBS washes and 1x D0 wash, then 200 μL parasites in complete D10H at a concentration of 1e6 parasites/mL were added. The plate was centrifuged to facilitate parasites-host cell contact (290 g, 5 min), and the plate was incubated at 37°C for 2 hours.

Stocks of inhibitors were all dissolved in DMSO; 200 mM benzyl-α-GalNAc (Cayman Chemical; 31747-100), 10 mM kifunensine (Sigma, K1140), and 10 mM eliglustat (Selleck Chemicals, S7852). For benzyl-α-GalNAc and kifunensine co-treatments, HFF cells were split from 12.5 cm^2^ flasks into 96 well plates (1 flask per plate), and grown for 3 days in presence of 2 mM BAG and 10 μM kifunensine (unless otherwise noted). For eliglustat and kifunensine co-treatments, HFF cells were split 1:9 and grown for 5 days in presence of the indicated drug, then seeded into 96 well plates. Rhoptry discharge assays were performed on day 7 after drug treatment. To confirm eliglustat affected sphingolipid levels, on day 5 HFF cells were stained in FACS buffer with Alexa Fluor 594 conjugated cholera toxin subunit B (CtxB-AF594, Invitrogen; C34777), which binds to the sphingolipid Gm1 [reference]. For rhoptry discharge assays, parasites in D10H at a concentration of 1e6 parasites / mL were added to host cells. The plate was centrifuged to facilitate parasites-host cell contact (290 g, 5 min), and the plate was incubated at 37°C for 2 hours.

After 2 hours at 37°C to allow for rhoptry discharge and nuclear translocation of phosphorylated STAT6, cells were washed once with PBS then fixed by addition of 100% ice-cold methanol for 8 minutes. Wells were then washed 3X with PBS to remove methanol, then blocked for 10 minutes in blocking buffer (PBS with 1% goat serum [Gibco; 16210072] and 5% IFS). Samples were stained with 1:10,000 guinea pig anti-*Tg*CDPK1 ^118^ and 1:1000 rabbit anti-pSTAT6 (HFF staining with Abcam ab188080, CHO staining with Cell Signaling #56554). Primary incubation was for 1 hour at RT or overnight at 4°C. Wells were washed 2X with PBS, then incubated in secondary staining solution (1:1000 anti-rabbit Alexa Fluor 488 [Life Technologies; A11034], 1:1000 anti-guinea pig Alexa Fluor 594, and 1:20,000 Hoescht 33258 [Santa Cruz Biotechnology; SC-39403]) for 1 h at RT. Wells were washed 3X with PBS then a final 200 μL PBS was added, and wells were imaged using a Cytation 3 or Cytation 5 Multimode reader (BioTek) using a 20X objective with laser autofocus. A custom Python script was used to identify nuclei in the Hoescht channel and extract nuclear pSTAT6 signal while masking out parasites using the anti-*Tg*CDPK1 counterstain. An R analysis script and GraphPad Prism were used to threshold the percentage of pSTAT6 positive nuclei and perform statistical analysis, respectively.

### Flow-based adhesion assays

HFF cells were grown to confluency in 0.8 mm height IbiTreat µ-Slide I Luer channel slides (ibidi). RH/Δ*ku80*/Δ*hxgprt*/dTomato (TgDV101) *T. gondii* parasites were syringe released and filtered with 5 μm pore filters to remove unlysed host cells and debris. Anti-SAG1 antibody (DG-52 monoclonal, ^133^) was preconjugated using AZDye 488 NHS Ester to make Anti-SAG1-488. The AZDye 488 reaction was quenched using 100 mM Tris and unconjugated fluorophore was removed using PD SpinTrap™ G-25 desalting and buffer-exchange columns (28918004, GE Healthcare/Cytiva), and the final concentration was estimated to be >0.8 mg/mL. *T. gondii* parasites were then centrifuged at 2000 g for 5 min, and resuspended in FluoroBrite DMEM (Gibco) containing 20 mM HEPES pH 7.4 (F0H) and 1:100 dilution of anti-SAG1-488. After 20-30 minutes, parasites were pelleted and resuspended in FluoroBrite DMEM containing 10% IFS and 20 mM HEPES (F10H). Parasites were diluted to 1×10^6^ parasites/mL in F10H containing PBS (vehicle) or 10 mM galactose, collected in a syringe and mounted in a syringe pump. The µ-Slide channel slide was mounted on a spinning disk confocal microscope with a dual camera set up and a 40X air objective (CSU-W1, Nikon) inside a temperature chamber set to 37°C. Silicone tubing with an inner diameter of 0.8 mm (10841, ibidi) was used to connect the syringe to the µ-Slide channel slide containing HFF cells, with the tubing passing through a bottle water bath inside the heated chamber. The syringe pump was set to a flow rate of 15 μL/min. Parasites were imaged for 30 minutes in dual camera mode (488/561 laser lines for AzDye488/dTomato), with 5 second intervals and a 9 plane Z-stack centred on the surface of the HFF cells. SAG1 staining was used to account for invasion events by Anti-SAG1 shedding ^21^. To analyze parasites under flow, maximum intensity projections were generated using NIS elements software, then Fiji ^128^ was used to analyze parasite tracks in the dTomato channel using the Trackmate Plugin (LoG spot detector and LAP tracker). Tracks present in the first 3 frames (15 seconds) or shorter than 3 frames were omitted from analysis. A rolling average of 3 concurrent frames was calculated to smooth out the parasite trajectories. Using Rstudio, parasites with a low initial rolling speed were also filtered out (<0.3 μm/second) to remove erroneous track switching from the LAP tracking algorithm. Triplicate experimental data was combined and hierarchical clustering was used to assess the distribution of log_2_(rolling minimum speed) for all data, and a cluster number of 6 was chosen for further analysis. The clusters were manually checked for appropriate cutoffs and it was observed that binning the top and bottom 3 clusters appropriately separated adhered and moving parasites. The displayed results were graphed and assessed using t-tests using R.

### Host cell perforation transient and invasion assays

Live calcium transient/invasion assays were carried out as previously described ^21^. Briefly, we used the protocol with Cal-520 AM as the calcium indicator, inclusion of 1 mM probenecid throughout the protocol, and live cell imaging solution containing 8 mM CaCl2 during live imaging. For each of the three replicate experiments, CHO cell lines (WT, SLC35A1 KO [Lec2], SLC35A2 KO [Lec8]) were each seeded in two chambers of an ibidi µ-Slide VI 0.4 (ibidi) in Ham’s F-12 medium with glutamine (Cytiva) containing 100 units/mL penicillin, and 100 µg/mL streptomycin, 10% heat-inactivated fetal bovine serum (Life Technologies), and incubated overnight at 37°C (with 5% CO2 and humidity) in preparation for live experiments. As a lower level of invasion was observed into CHO cells relative to HFF cells, 2400 high quality frames (192 seconds) were processed from each capture compared to the published protocol ^21^. Due to high levels of baseline intracellular calcium activity in the CHO cells, automated identification of calcium transients was confirmed by visualization of each event as a change in fluorescence over time plot.

### Glycan bead affinity purifications

*T. gondii* RH/Δ*ku80/*Δ*hxgprt/*dTomato (TgDV101) parasites were grown in 15 cm^2^ plates of confluent HFF cells for 2 days. Parasites were harvested, filtered with 5 μm pore filters to remove host cells and debris, washed 2X with PBS via pelleting (5 min, 2000 g) and resuspension. Parasites were pelleted again, the final PBS wash was removed and the pellet was flash frozen in liquid N_2_. Pellets were lysed on ice by thawing directly into cold SugarAP buffer (PBS with 1 mM CaCl_2_, MgCl_2_, 0.5% Igepal CA-630, 1 mM benzonase [Millipore; E1014-25KU], and 1X protease inhibitor cocktail [Thermo; 87786]). While parasites were lysing, 200 μL bead slurry (100 μL final bed) of D-mannose agarose (Sigma; M6400), D-galactose agarose (Pierce; PI20372), and α-lactose agarose were added to spin columns (Pierce; 89879) and prepared by washing with 10 gel-bed volumes (4×250 μL) of SugarAP buffer; 40 g spins were used throughout for all steps. The parasite lysate was clarified by a centrifugation to remove insoluble debris (4°C, 11000 g, 10 min), and the soluble lysate flowed over the resins. Resins were then washed for 20 gel-bed volumes (8x 250 μL) using SugarAP buffer. Bound proteins were then specifically eluted by 100 mM of the free cognate sugar in SugarAP buffer (i.e. mannose, galactose, lactose), using 7.5 gel-bed volumes (3x 250 μL).

Eluted proteins were prepared for mass spectrometry by SP3 bead digestion and clean-up as previously described ^10,134^. Briefly, proteins were reduced with TCEP (Pierce, PI20490) and alkylated with chloroacetamide (Thermo, 148415000), precipitated onto hydrophilic and hydrophobic SP3 beads using ethanol, washed in 80% ethanol, and digested overnight with Trypsin/LysC mix (Thermo, A40007). Eluted peptides were lyophilized and directly injected for DDA mass spectrometry on a Thermo Exploris 480 mass spectrometer and analyzed using Proteome Discoverer 2.4 (Thermo).

## ACKNOWLEDGEMENTS

We would like to thank Monty Krieger, Pamela Stanley, and Richard Cummings for the CHO cell mutants; MyLJHang Huynh and Vern B. Carruthers for the *T. gondii* Δ*spatr* strain; and L. David Sibley for the MIC2 and SAG1 antibodies. We are also grateful to Maryse Lebrun for helpful advice and discussions, to the VEuPathDB team for maintaining crucial resources used in the preparation of this manuscript, and to the Whitehead Quantitative Proteomics Core (QPC) and the Function Genomics Platform (FGP) for experimental assistance. This project was supported by NIH grants (AI144369 and AI158501) and by the Burroughs Wellcome Fund (grant 1021330) awarded to SL, NIH grant AI169067 to GEW, and a Long-term Fellowship from the Human Frontier Science Program to DV (LT000890/2021LJL).

---

**Figure S1.**
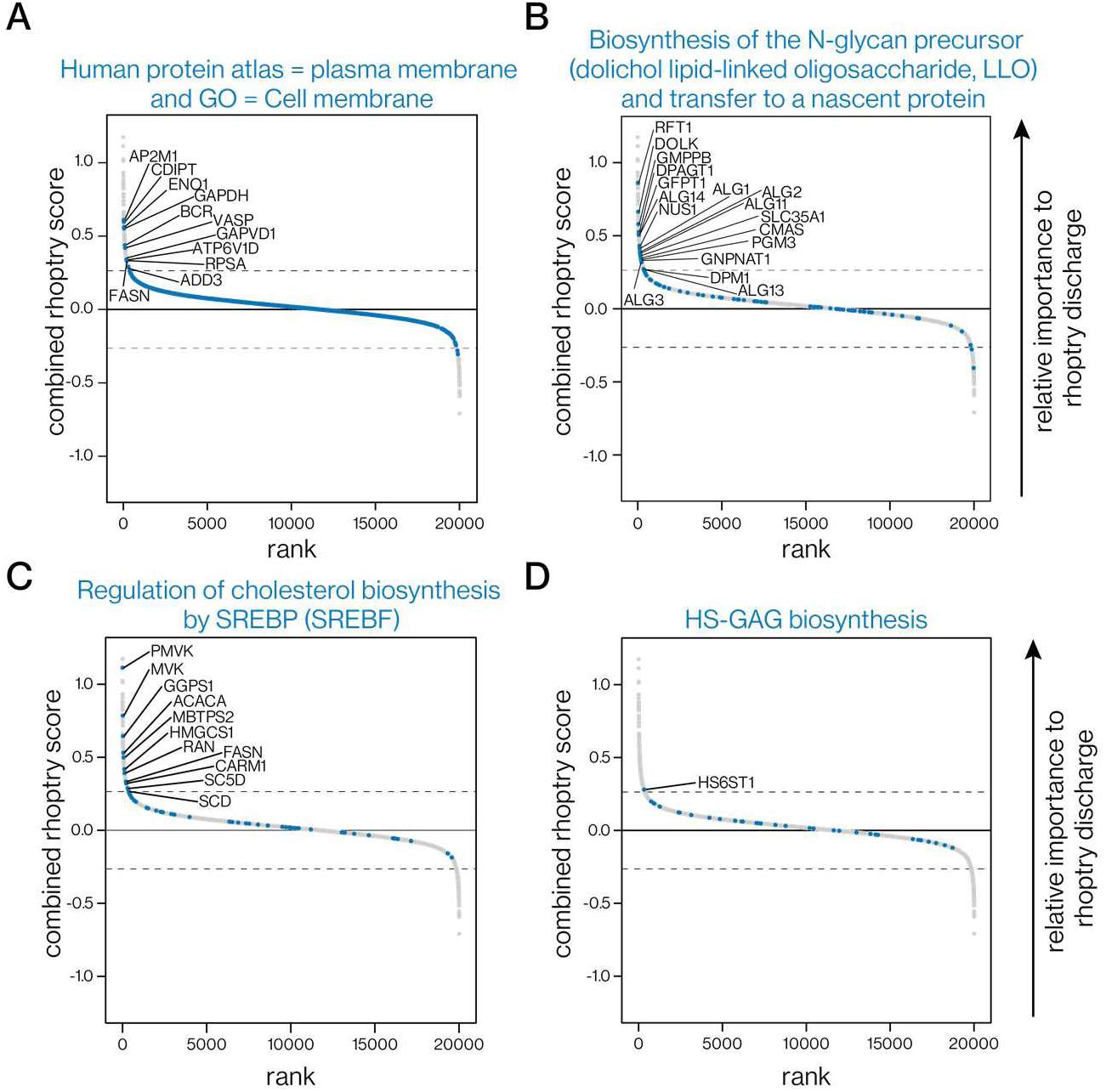
Involvement of relevant gene categories in rhoptry discharge. **(A)** Rank ordered position of genes that are displayed in the plasma membrane or surface exposed as annotated by GO terms and the human protein atlas in the rhoptry discharge screen. Dashed line indicates MLE estimation of s.d. = 3. All genes above this threshold are labeled. **(B)** Rank ordered position of genes required for N-glycan dolichol precursor biosynthesis and transfer to a nascent protein in the rhoptry discharge screen. Genes displayed are from the Reactome pathway R-HSA-446193. Dashed line indicates MLE estimation of s.d = 3. All genes above this threshold are labeled. **(C)** Rank ordered position of genes required for regulation of cholesterol biosynthesis in the rhoptry discharge screen. Genes displayed are from the Reactome pathway R-HSA-1655829. Dashed line indicates MLE estimation of s.d. = 3. All genes above this threshold are labeled. **(D)** Rank ordered position of genes in the heparin sulfate GAG biosynthesis pathway in the rhoptry discharge screen. Genes displayed are from the Reactome pathway R-HSA-2022928. Dashed line indicates MLE estimation of s.d = 3. All genes above this threshold are labeled.

**Figure S2.**
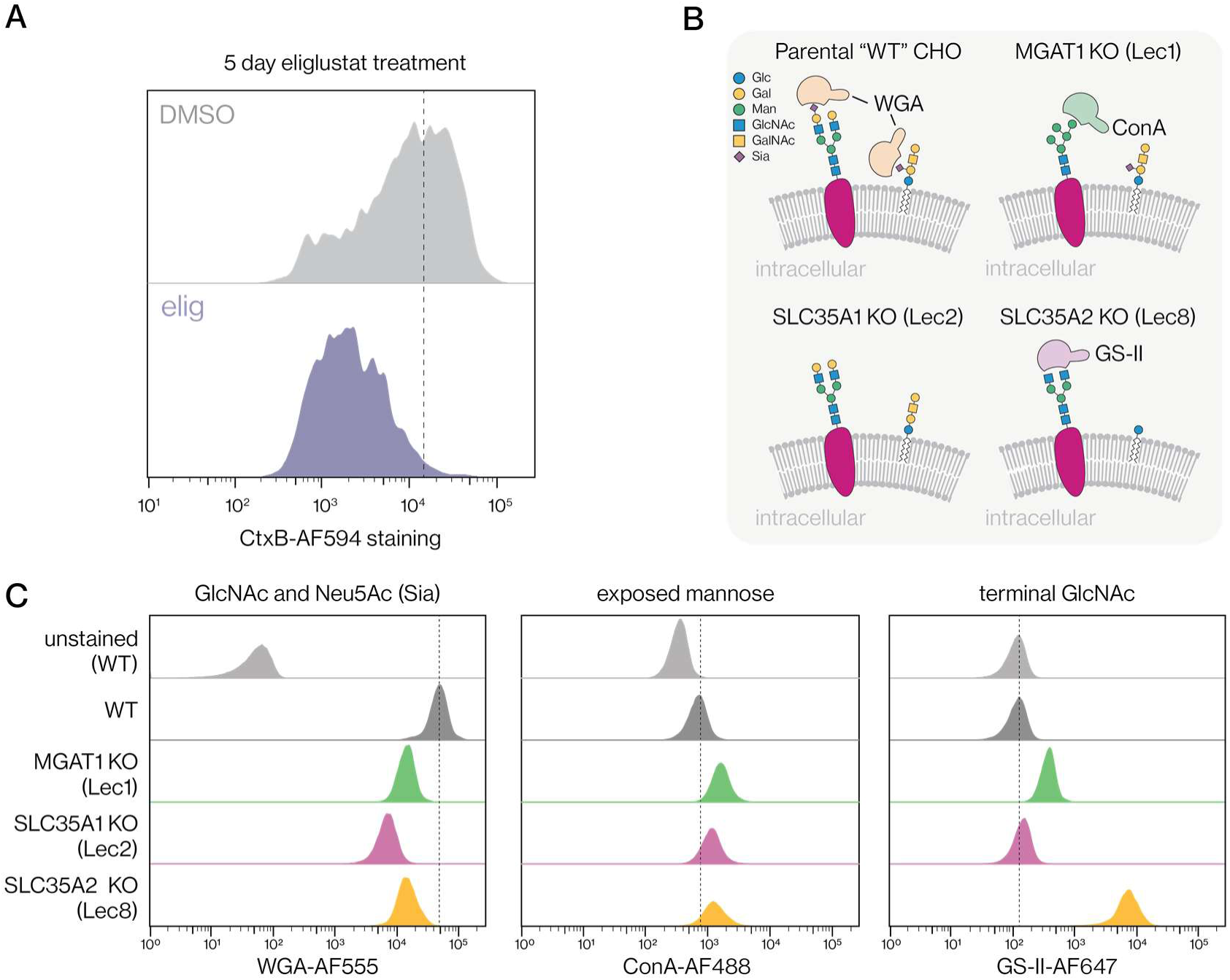
Host cell drug treatments and knockouts result in expected surface glycosylation patterns. **(A)** Flow cytometry of plasma membrane glycosphingolipid Gm1 content after treating human fibroblasts with vehicle control or the UGCG inhibitor eliglustat for 5 days, as assessed Alexa Fluor 594-conjugated Cholera toxin B domain (CtxB-AF594). **(B)** The collected CHO cells and their expected glycosylation patterns. **(C)** Verification of CHO cell mutants surface glycosylation profiles by lectin-labeling and flow cytometry. CHO cells (WT [Pro5-parental], MGAT1 KO [Lec1], SLC35A1 KO [Lec2], and SLC35A2 KO [Lec8]) were stained with GlcNAc and Neu5Ac (sialic acid) binding WGA (Alexa Fluor 555 conjugated), mannose-binding ConA (Alexa Fluor 488 conjugated), and terminal GlcNAc-binding GS-II (Alexa Fluor 647 conjugated).

**Figure S3.**
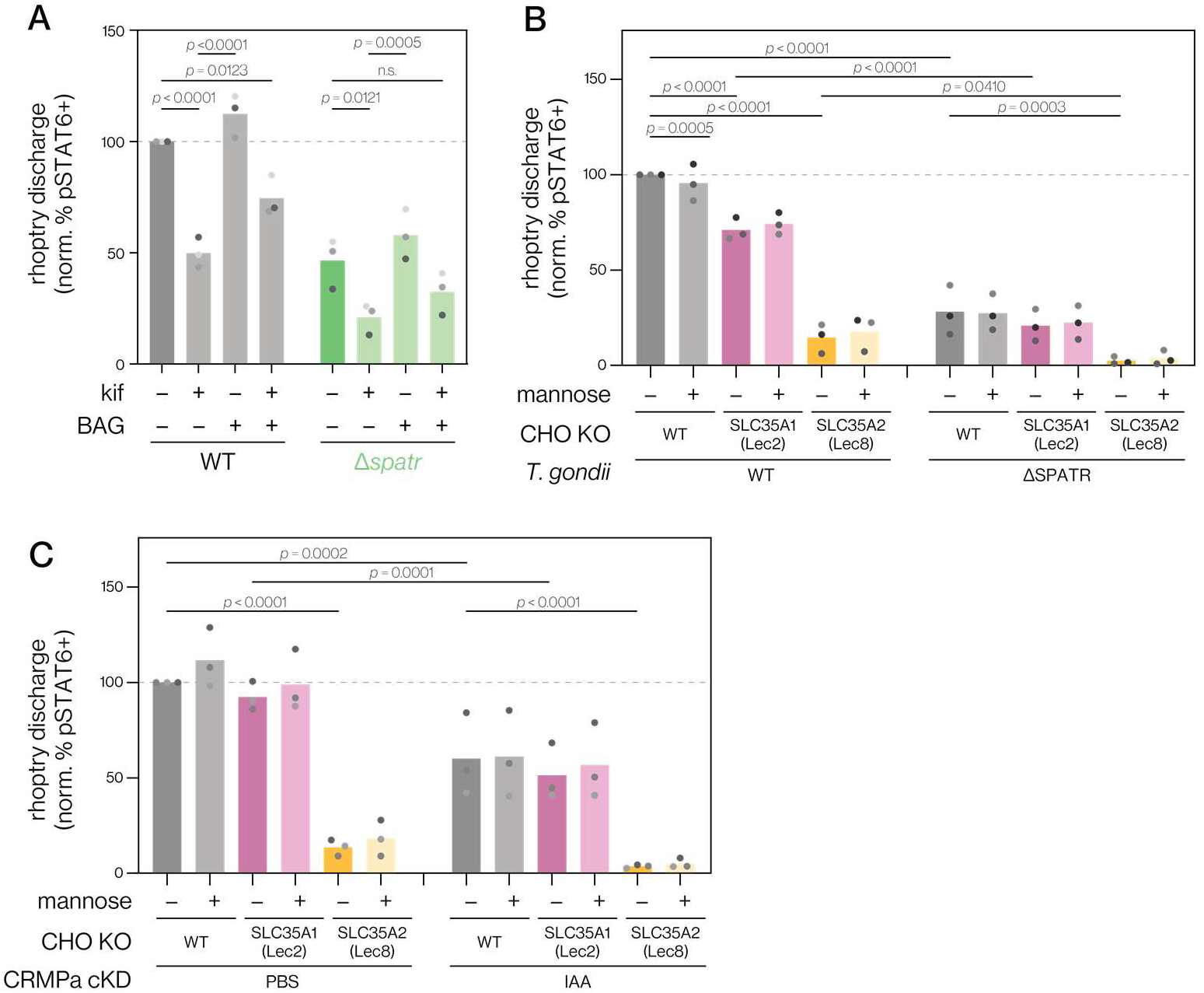
Supporting data demonstrating that CLAMP and CRMP complexes additively interact with galactose and N-glycans. **(A)** The interaction of the CLAMP complex with N-glycans and O-glycans in rhoptry discharge was assessed by treating human fibroblasts for 3 days with kif (10 μM) or BAG (2 mM) then incubating with Δ*spatr* or wild-type parental parasites. Rhoptry discharge was assessed by the relative percent of pSTAT6–positive nuclei. Mean of *n* = 3 biological replicates is plotted; *p*-values were derived from two-way ANOVA with Tukey’s post hoc multiple comparison test. Note the values for the wild-type parasites condition are the same data as those in Figure 2E. **(B-C)** Evaluation of the interaction of CHO galactose (SLC35A2 KO) and sialic acid (SLC35A1 KO) knockouts combined with mannose addition against the (**A**) CLAMP and (**B**) CRMP complexes in rhoptry discharge. Host cells were challenged with the indicated CLAMP perturbation (wild-type or Δ*spatr* parasites, **A**) or CRMP perturbation (CRMPa cKD treated with IAA or PBS vehicle, **B**) in the presence of 10 mM of mannose, then rhoptry discharge was assessed by the relative percent of pSTAT6–positive nuclei. Mean of *n* = 3 biological replicates is plotted; *p*-values are derived from two-way ANOVA testing with multiple comparison FDR correction using the Benjamini, Krieger, and Yekutieli two-stage step-up method. For both **B** and **C**, data was analyzed as part of the same datasets as Figure 3C and **3D**, respectively.

**Figure S4.**
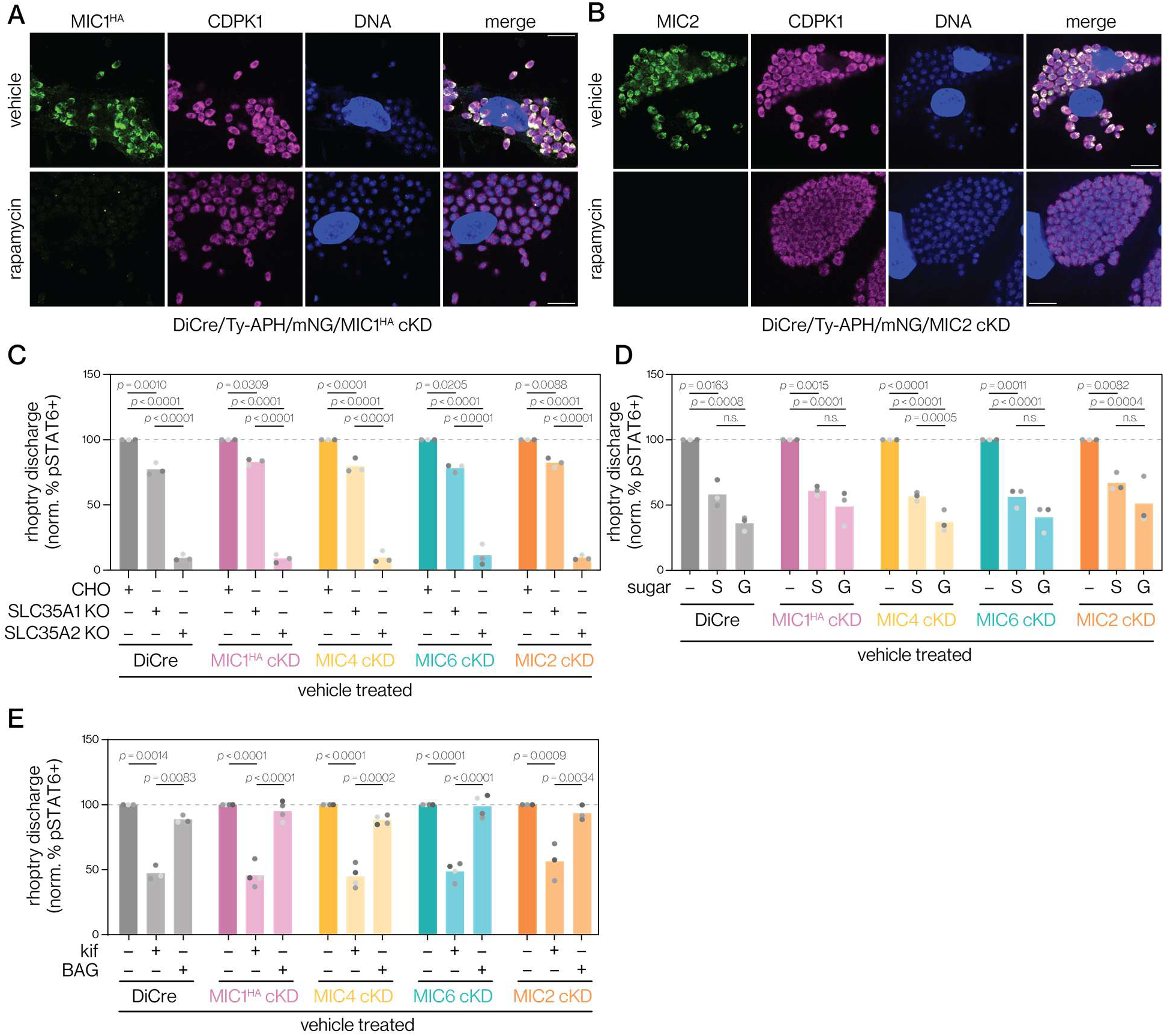
Strain validation and vehicle-treated conditions for MIC1/4/6 and MIC2 interactions with glycans in rhoptry discharge. **(A-B)** Evaluation of MIC1^HA^ **(A)** and MIC2 **(B)** microneme localization following rapamycin treatment of cKD strains. Parasites were treated for 2 h with rapamycin, then allowed to grow intracellularly for 48 h prior to fixation and staining. anti-CDPK1 was used as a *T. gondii* cytosolic control and Hoechst 33258 used to stain DNA. **(C)** Challenge of vehicle-treated DiCre, MIC1^HA^, MIC4, MIC6, and MIC2 cKD strains against WT, SLC35A1 KO, and SLC35A2 KO CHO cells. Control pSTAT6 assay conditions for rapamycin-induced knockdowns in Figure 4C. Mean of *n* = 3 biological replicates is plotted; *p*-values derived from analysis of each strain separately (vehicle and knockdown conditions) using two-way ANOVA with Tukey’s post hoc multiple comparison test. **(D)** Challenge of vehicle-treated DiCre MIC1^HA^, MIC4, MIC6, and MIC2 cKD strains against human fibroblast in presence of 10 mM of galactose (“g”), sialic acid (“s”) or vehicle (“-”). Control pSTAT6 assay conditions for rapamycin-induced knockdowns in Figure 4D. Mean of *n* = 3 biological replicates is plotted; *p*-values derived from analysis of each strain separately (vehicle and knockdown conditions) using two-way ANOVA with Tukey’s post hoc multiple comparison test. **(E)** Challenge of vehicle-treated treatment DiCre, MIC1^HA^, MIC4, MIC6, and MIC2 cKD strains against human fibroblasts treated for 4 days with kif (10 μM) or BAG (2 mM). Control pSTAT6 assay conditions for rapamycin-induced knockdowns in Figure 4E. Mean of *n* = 3 (DiCre and MIC2 cKD) or *n = 4* (MIC1, MIC4, MIC6 cKDs) biological replicates is plotted; *p*-values derived from analysis of each strain separately (vehicle and knockdown conditions) using two-way ANOVA with Tukey’s post hoc multiple comparison test.

**Figure S5.**
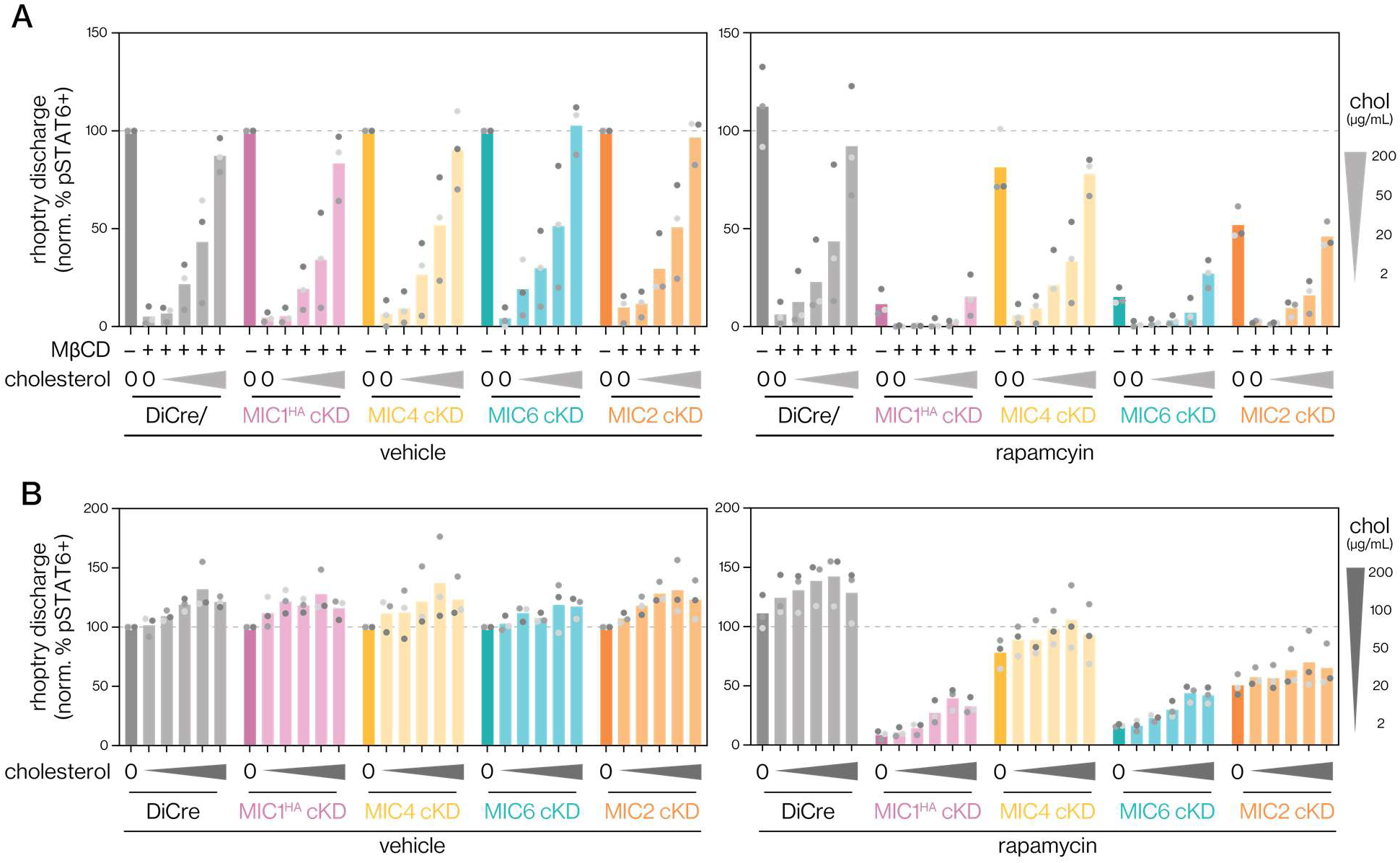
Underlying data for cholesterol level modulation interactions with the MIC1/4/6 complex. **(A)** Non log scaled data underlying cholesterol removal and reconstitution experiments in Figure 5B-F. Cholesterol reconstitution levels are displayed in the legend on right. **(B)** Non log scaled data underlying increased cholesterol experiments Figure 5B-F. Excess added cholesterol levels are displayed in the legend on right.

## TABLE LEGENDS

**Table S1. MAGeCK MLE analysis of the host-directed rhoptry dependency screen**

**Table S2. DDA-MS analysis of glycan bead precipitation.**

## REFERENCES

1. Weiss, G.E., Gilson, P.R., Taechalertpaisarn, T., Tham, W.-H., de Jong, N.W.M., Harvey, K.L., Fowkes, F.J.I., Barlow, P.N., Rayner, J.C., Wright, G.J., et al. (2015). Revealing the sequence and resulting cellular morphology of receptor-ligand interactions during Plasmodium falciparum invasion of erythrocytes. PLoS Pathog. 11, e1004670.

2. Wetzel, D.M., Schmidt, J., Kuhlenschmidt, M.S., Dubey, J.P., and Sibley, L.D. (2005). Gliding motility leads to active cellular invasion by Cryptosporidium parvum sporozoites. Infect. Immun. 73, 5379–5387.

3. Carruthers, V.B., and Boothroyd, J.C. (2007). Pulling together: an integrated model of Toxoplasma cell invasion. Curr. Opin. Microbiol. 10, 83–89.

4. Bichet, M., Touquet, B., Gonzalez, V., Florent, I., Meissner, M., and Tardieux, I. (2016). Genetic impairment of parasite myosin motors uncovers the contribution of host cell membrane dynamics to Toxoplasma invasion forces. BMC Biol. 14, 97.

5. Carruthers, V.B., and Sibley, L.D. (1997). Sequential protein secretion from three distinct organelles of Toxoplasma gondii accompanies invasion of human fibroblasts. Eur. J. Cell Biol. 73, 114–123.

6. Bisio, H., and Soldati-Favre, D. (2019). Signaling cascades governing entry into and exit from host cells by Toxoplasma gondii. Annu. Rev. Microbiol. 73, 579–599.

7. Carruthers, V.B., and Sibley, L.D. (1999). Mobilization of intracellular calcium stimulates microneme discharge in Toxoplasma gondii. Mol. Microbiol. 31, 421–428.

8. Cova, M.M., Lamarque, M.H., and Lebrun, M. (2022). How Apicomplexa Parasites Secrete and Build Their Invasion Machinery. Annu. Rev. Microbiol. 76, 619–640.

9. Cowman, A.F., Tonkin, C.J., Tham, W.-H., and Duraisingh, M.T. (2017). The molecular basis of erythrocyte invasion by malaria parasites. Cell Host Microbe 22, 232–245.

10. Valleau, D., Sidik, S.M., Godoy, L.C., Acevedo-Sánchez, Y., Pasaje, C.F.A., Huynh, M.-H., Carruthers, V.B., Niles, J.C., and Lourido, S. (2023). A conserved complex of microneme proteins mediates rhoptry discharge in Toxoplasma. EMBO J. 42, e113155.

11. Singer, M., Simon, K., Forné, I., and Meissner, M. (2023). A central CRMP complex essential for invasion in Toxoplasma gondii. PLoS Biol. 21, e3001937.

12. Sparvoli, D., Delabre, J., Penarete-Vargas, D.M., Kumar Mageswaran, S., Tsypin, L.M., Heckendorn, J., Theveny, L., Maynadier, M., Mendonça Cova, M., Berry-Sterkers, L., et al. (2022). An apical membrane complex for triggering rhoptry exocytosis and invasion in Toxoplasma. EMBO J. 41, e111158.

13. Kessler, H., Herm-Götz, A., Hegge, S., Rauch, M., Soldati-Favre, D., Frischknecht, F., and Meissner, M. (2008). Microneme protein 8--a new essential invasion factor in Toxoplasma gondii. J. Cell Sci. 121, 947–956.

14. Pace, D.A., McKnight, C.A., Liu, J., Jimenez, V., and Moreno, S.N.J. (2014). Calcium entry in Toxoplasma gondii and its enhancing effect of invasion-linked traits. J. Biol. Chem. 289, 19637–19647.

15. Brown, K.M., Lourido, S., and Sibley, L.D. (2016). Serum albumin stimulates protein kinase G-dependent microneme secretion in Toxoplasma gondii. J. Biol. Chem. 291, 9554–9565.

16. Tagoe, D.N.A., Drozda, A.A., Falco, J.A., Bechtel, T.J., Weerapana, E., and Gubbels, M.-J. (2021). Ferlins and TgDOC2 in Toxoplasma Microneme, Rhoptry and Dense Granule Secretion. Life 11. 10.3390/life11030217.

17. Farrell, A., Thirugnanam, S., Lorestani, A., Dvorin, J.D., Eidell, K.P., Ferguson, D.J.P., Anderson-White, B.R., Duraisingh, M.T., Marth, G.T., and Gubbels, M.-J. (2012). A DOC2 Protein Identified by Mutational Profiling Is Essential for Apicomplexan Parasite Exocytosis. Science 335, 218–221.

18. Frénal, K., Dubremetz, J.-F., Lebrun, M., and Soldati-Favre, D. (2017). Gliding motility powers invasion and egress in Apicomplexa. Nat. Rev. Microbiol. 15, 645–660.

19. Graindorge, A., Frénal, K., Jacot, D., Salamun, J., Marq, J.B., and Soldati-Favre, D. (2016). The Conoid Associated Motor MyoH Is Indispensable for Toxoplasma gondii Entry and Exit from Host Cells. PLoS Pathog. 12, e1005388.

20. Lamarque, M.H., Roques, M., Kong-Hap, M., Tonkin, M.L., Rugarabamu, G., Marq, J.-B., Penarete-Vargas, D.M., Boulanger, M.J., Soldati-Favre, D., and Lebrun, M. (2014). Plasticity and redundancy among AMA-RON pairs ensure host cell entry of Toxoplasma parasites. Nat. Commun. 5, 4098.

21. Male, F., Kegawa, Y., Blank, P.S., Jiménez-Munguía, I., Sidik, S.M., Valleau, D., Lourido, S., Lebrun, M., Zimmerberg, J., and Ward, G.E. (2025). Perforation of the host cell plasma membrane during Toxoplasma invasion requires rhoptry exocytosis. EMBO Rep. 10.1038/s44319-025-00564-9.

22. Huynh, M.-H., and Carruthers, V.B. (2006). Toxoplasma MIC2 is a major determinant of invasion and virulence. PLoS Pathog. 2, e84.

23. Tyler, J.S., and Boothroyd, J.C. (2011). The C-terminus of Toxoplasma RON2 provides the crucial link between AMA1 and the host-associated invasion complex. PLoS Pathog. 7, e1001282.

24. Lamarque, M., Besteiro, S., Papoin, J., Roques, M., Vulliez-Le Normand, B., Morlon-Guyot, J., Dubremetz, J.-F., Fauquenoy, S., Tomavo, S., Faber, B.W., et al. (2011). The RON2-AMA1 interaction is a critical step in moving junction-dependent invasion by apicomplexan parasites. PLoS Pathog. 7, e1001276.

25. Broncel, M., Dominicus, C., Vigetti, L., Nofal, S.D., Bartlett, E.J., Touquet, B., Hunt, A., Wallbank, B.A., Federico, S., Matthews, S., et al. (2020). Profiling of myristoylation in Toxoplasma gondii reveals an N-myristoylated protein important for host cell penetration. Elife 9, e57861.

26. Mageswaran, S.K., Guérin, A., Theveny, L.M., Chen, W.D., Martinez, M., Lebrun, M., Striepen, B., and Chang, Y.-W. (2021). In situ ultrastructures of two evolutionarily distant apicomplexan rhoptry secretion systems. Nat. Commun. 12, 4983.

27. Martinez, M., Chen, W.D., Cova, M.M., Molnár, P., Mageswaran, S.K., Guérin, A., John, A.R.O., Lebrun, M., and Chang, Y.-W. (2022). Rhoptry secretion system structure and priming in Plasmodium falciparum revealed using in situ cryo-electron tomography. Nat Microbiol. 10.1038/s41564-022-01171-3.

28. Segev-Zarko, L.-A., Dahlberg, P.D., Sun, S.Y., Pelt, D.M., Kim, C.Y., Egan, E.S., Sethian, J.A., Chiu, W., and Boothroyd, J.C. (2022). Cryo-electron tomography with mixed-scale dense neural networks reveals key steps in deployment of Toxoplasma invasion machinery. PNAS Nexus 1. 10.1093/pnasnexus/pgac183.

29. Kegawa, Y., Male, F., Jiménez-Munguía, I., Blank, P.S., Mekhedov, E., Ward, G.E., and Zimmerberg, J. (2025). The invasion pore induced by Toxoplasma gondii. EMBO Rep. 10.1038/s44319-025-00565-8.

30. Geoghegan, N.D., Evelyn, C., Whitehead, L.W., Pasternak, M., McDonald, P., Triglia, T., Marapana, D.S., Kempe, D., Thompson, J.K., Mlodzianoski, M.J., et al. (2021). 4D analysis of malaria parasite invasion offers insights into erythrocyte membrane remodeling and parasitophorous vacuole formation. Nat. Commun. 12, 3620.

31. Ong, Y.-C., Reese, M.L., and Boothroyd, J.C. (2010). Toxoplasma rhoptry protein 16 (ROP16) subverts host function by direct tyrosine phosphorylation of STAT6. J. Biol. Chem. 285, 28731–28740.

32. Koshy, A.A., Fouts, A.E., Lodoen, M.B., Alkan, O., Blau, H.M., and Boothroyd, J.C. (2010). Toxoplasma secreting Cre recombinase for analysis of host-parasite interactions. Nat. Methods 7, 307–309.

33. Koshy, A.A., Dietrich, H.K., Christian, D.A., Melehani, J.H., Shastri, A.J., Hunter, C.A., and Boothroyd, J.C. (2012). Toxoplasma co-opts host cells it does not invade. PLoS Pathog. 8, e1002825.

34. Lodoen, M.B., Gerke, C., and Boothroyd, J.C. (2010). A highly sensitive FRET-based approach reveals secretion of the actin-binding protein toxofilin during Toxoplasma gondii infection. Cell. Microbiol. 12, 55–66.

35. Harper, J.M., Hoff, E.F., and Carruthers, V.B. (2004). Multimerization of the Toxoplasma gondii MIC2 integrin-like A-domain is required for binding to heparin and human cells. Mol. Biochem. Parasitol. 134, 201–212.

36. Gras, S., Jackson, A., Woods, S., Pall, G., Whitelaw, J., Leung, J.M., Ward, G.E., Roberts, C.W., and Meissner, M. (2017). Parasites lacking the micronemal protein MIC2 are deficient in surface attachment and host cell egress, but remain virulent in vivo. Wellcome Open Res. 2, 32.

37. Carruthers, V.B., Håkansson, S., Giddings, O.K., and Sibley, L.D. (2000). Toxoplasma gondii uses sulfated proteoglycans for substrate and host cell attachment. Infect Immun 68, 4005–4011.

38. Coppens, I., and Joiner, K.A. (2003). Host but not parasite cholesterol controls Toxoplasma cell entry by modulating organelle discharge. Mol. Biol. Cell 14, 3804–3820.

39. Tahara, M., Andrabi, S.B.A., Matsubara, R., Aonuma, H., and Nagamune, K. (2016). A host cell membrane microdomain is a critical factor for organelle discharge by Toxoplasma gondii. Parasitol. Int. 65, 378–388.

40. Farrell, B., Alam, N., Hart, M.N., Jamwal, A., Ragotte, R.J., Walters-Morgan, H., Draper, S.J., Knuepfer, E., and Higgins, M.K. (2024). The PfRCR complex bridges malaria parasite and erythrocyte during invasion. Nature 625, 578–584.

41. Day, C.J., Favuzza, P., Bielfeld, S., Haselhorst, T., Seefeldt, L., Hauser, J., Shewell, L.K., Flueck, C., Poole, J., Jen, F.E.-C., et al. (2024). The essential malaria protein PfCyRPA targets glycans to invade erythrocytes. Cell Rep. 43, 114012.

42. Dos Santos Pacheco, N., Brusini, L., Haase, R., Tosetti, N., Maco, B., Brochet, M., Vadas, O., and Soldati-Favre, D. (2022). Conoid extrusion regulates glideosome assembly to control motility and invasion in Apicomplexa. Nat. Microbiol. 7, 1777–1790.

43. Hunt, A., Russell, M.R.G., Wagener, J., Kent, R., Carmeille, R., Peddie, C.J., Collinson, L., Heaslip, A., Ward, G.E., and Treeck, M. (2019). Differential requirements for cyclase-associated protein (CAP) in actin-dependent processes of Toxoplasma gondii. Elife 8. 10.7554/eLife.50598.

44. Pieperhoff, M.S., Pall, G.S., Jiménez-Ruiz, E., Das, S., Melatti, C., Gow, M., Wong, E.H., Heng, J., Müller, S., Blackman, M.J., et al. (2015). Conditional U1 Gene Silencing in Toxoplasma gondii. PLoS One 10. 10.1371/journal.pone.0130356.

45. Inglis, A.J., Guna, A., Gálvez-Merchán, Á., Pal, A., Esantsi, T.K., Keys, H.R., Frenkel, E.M., Oania, R., Weissman, J.S., and Voorhees, R.M. (2023). Coupled protein quality control during nonsense-mediated mRNA decay. J. Cell Sci. 136, jcs261216.

46. Ashburner, M., Ball, C.A., Blake, J.A., Botstein, D., Butler, H., Cherry, J.M., Davis, A.P., Dolinski, K., Dwight, S.S., Eppig, J.T., et al. (2000). Gene ontology: tool for the unification of biology. The Gene Ontology Consortium. Nat. Genet. 25, 25–29.

47. Gene Ontology Consortium, Aleksander, S.A., Balhoff, J., Carbon, S., Cherry, J.M., Drabkin, H.J., Ebert, D., Feuermann, M., Gaudet, P., Harris, N.L., et al. (2023). The Gene Ontology knowledgebase in 2023. Genetics 224, iyad031.

48. Thul, P.J., Åkesson, L., Wiking, M., Mahdessian, D., Geladaki, A., Ait Blal, H., Alm, T., Asplund, A., Björk, L., Breckels, L.M., et al. (2017). A subcellular map of the human proteome. Science 356. 10.1126/science.aal3321.

49. Milacic, M., Beavers, D., Conley, P., Gong, C., Gillespie, M., Griss, J., Haw, R., Jassal, B., Matthews, L., May, B., et al. (2024). The reactome pathway knowledgebase 2024. Nucleic Acids Res. 52, D672–D678.

50. Sukhanova, A., Gorin, A., Serebriiskii, I.G., Gabitova, L., Zheng, H., Restifo, D., Egleston, B.L., Cunningham, D., Bagnyukova, T., Liu, H., et al. (2013). Targeting C4-demethylating genes in the cholesterol pathway sensitizes cancer cells to EGF receptor inhibitors via increased EGF receptor degradation. Cancer Discov. 3, 96–111.

51. Fidelito, G., Todorovski, I., Cluse, L., Vervoort, S.J., Taylor, R.A., and Watt, M.J. (2025). Lipid-metabolism-focused CRISPR screens identify enzymes of the mevalonate pathway as essential for prostate cancer growth. Cell Rep. 44, 115470.

52. Schjoldager, K.T., Narimatsu, Y., Joshi, H.J., and Clausen, H. (2020). Global view of human protein glycosylation pathways and functions. Nat. Rev. Mol. Cell Biol. 21, 729–749.

53. Mullen, P.J., Yu, R., Longo, J., Archer, M.C., and Penn, L.Z. (2016). The interplay between cell signalling and the mevalonate pathway in cancer. Nat. Rev. Cancer 16, 718–731.

54. Besteiro, S., Bertrand-Michel, J., Lebrun, M., Vial, H., and Dubremetz, J.-F. (2008). Lipidomic analysis of Toxoplasma gondii tachyzoites rhoptries: further insights into the role of cholesterol. Biochem. J. 415, 87–96.

55. Fan, Y.-M., Zhang, Q.-Q., Pan, M., Hou, Z.-F., Fu, L., Xu, X., and Huang, S.-Y. (2024). Toxoplasma gondii sustains survival by regulating cholesterol biosynthesis and uptake via SREBP2 activation. J. Lipid Res. 65, 100684.

56. Håkansson, S., Charron, A.J., and Sibley, L.D. (2001). Toxoplasma evacuoles: a two-step process of secretion and fusion forms the parasitophorous vacuole. EMBO J. 20, 3132– 3144.

57. Subczynski, W.K., Pasenkiewicz-Gierula, M., Widomska, J., Mainali, L., and Raguz, M. (2017). High cholesterol/low cholesterol: Effects in biological membranes: A review. Cell Biochem. Biophys. 75, 369–385.

58. Huynh, M.-H., Boulanger, M.J., and Carruthers, V.B. (2014). A Conserved Apicomplexan Microneme Protein Contributes to Toxoplasma gondii Invasion and Virulence. Infect. Immun. 82, 4358–4368.

59. Mahammad, S., and Parmryd, I. (2015). Cholesterol depletion using methyl-β-cyclodextrin. Methods Mol. Biol. 1232, 91–102.

60. Wolfe, A.L., Zhou, Q., Toska, E., Galeas, J., Ku, A.A., Koche, R.P., Bandyopadhyay, S., Scaltriti, M., Lebrilla, C.B., McCormick, F., et al. (2021). UDP-glucose pyrophosphorylase 2, a regulator of glycogen synthesis and glycosylation, is critical for pancreatic cancer growth. Proc. Natl. Acad. Sci. U. S. A. 118, e2103592118.

61. Wang, Y., Maeda, Y., Liu, Y.-S., Takada, Y., Ninomiya, A., Hirata, T., Fujita, M., Murakami, Y., and Kinoshita, T. (2020). Cross-talks of glycosylphosphatidylinositol biosynthesis with glycosphingolipid biosynthesis and ER-associated degradation. Nat. Commun. 11, 860.

62. Drabavicius, G., and Daelemans, D. (2021). Intermedilysin cytolytic activity depends on heparan sulfates and membrane composition. PLoS Genet. 17, e1009387.

63. Kim, S., Wolfe, A., and Kim, S.E. (2021). Targeting cancer’s sweet spot: UGP2 as a therapeutic vulnerability. Mol. Cell. Oncol. 8, 1990676.

64. Führing, J.I., Cramer, J.T., Schneider, J., Baruch, P., Gerardy-Schahn, R., and Fedorov, R. (2015). A quaternary mechanism enables the complex biological functions of octameric human UDP-glucose pyrophosphorylase, a key enzyme in cell metabolism. Sci. Rep. 5, 9618.

65. Elbein, A.D., Tropea, J.E., Mitchell, M., and Kaushal, G.P. (1990). Kifunensine, a potent inhibitor of the glycoprotein processing mannosidase I. J. Biol. Chem. 265, 15599–15605.

66. Chang, V.T., Crispin, M., Aricescu, A.R., Harvey, D.J., Nettleship, J.E., Fennelly, J.A., Yu, C., Boles, K.S., Evans, E.J., Stuart, D.I., et al. (2007). Glycoprotein structural genomics: solving the glycosylation problem. Structure 15, 267–273.

67. Wang, S.-S., Solar, V.D., Yu, X., Antonopoulos, A., Friedman, A.E., Agarwal, K., Garg, M., Ahmed, S.M., Addhya, A., Nasirikenari, M., et al. (2021). Efficient inhibition of O-glycan biosynthesis using the hexosamine analog Ac5GalNTGc. Cell Chem. Biol. 28, 699–710.e5.

68. Kuan, S.F., Byrd, J.C., Basbaum, C., and Kim, Y.S. (1989). Inhibition of mucin glycosylation by aryl-N-acetyl-α-galactosaminides in human colon cancer cells. J. Biol. Chem. 264, 19271–19277.

69. Kong, W.-Z., and Fujita, M. (2025). GlycoMaple: recent updates and applications in visualization and analysis of glycosylation pathways. Anal Bioanal Chem 417, 885–894.

70. McDonald, A.G., Hayes, J.M., and Davey, G.P. (2016). Metabolic flux control in glycosylation. Curr. Opin. Struct. Biol. 40, 97–103.

71. Scheper, A.F., Schofield, J., Bohara, R., Ritter, T., and Pandit, A. (2023). Understanding glycosylation: Regulation through the metabolic flux of precursor pathways. Biotechnol. Adv. 67, 108184.

72. Dong, L., Cao, Z., Chen, M., Liu, Y., Ma, X., Lu, Y., Zhang, Y., Feng, K., Zhang, Y., Meng, Z., et al. (2024). Inhibition of glycosphingolipid synthesis with eliglustat in combination with immune checkpoint inhibitors in advanced cancers: preclinical evidence and phase I clinical trial. Nat Commun 15, 6970.

73. Stirnemann, J., Belmatoug, N., Camou, F., Serratrice, C., Froissart, R., Caillaud, C., Levade, T., Astudillo, L., Serratrice, J., Brassier, A., et al. (2017). A review of Gaucher disease pathophysiology, clinical presentation and treatments. Int. J. Mol. Sci. 18, 441.

74. Poole, J., Day, C.J., von Itzstein, M., Paton, J.C., and Jennings, M.P. (2018). Glycointeractions in bacterial pathogenesis. Nat. Rev. Microbiol. 16, 440–452.

75. Bucior, I., Abbott, J., Song, Y., Matthay, M.A., and Engel, J.N. (2013). Sugar administration is an effective adjunctive therapy in the treatment of Pseudomonas aeruginosa pneumonia. Am. J. Physiol. Lung Cell. Mol. Physiol. 305, L352–L363.

76. Friedrich, N., Santos, J.M., Liu, Y., Palma, A.S., Leon, E., Saouros, S., Kiso, M., Blackman, M.J., Matthews, S., Feizi, T., et al. (2010). Members of a novel protein family containing microneme adhesive repeat domains act as sialic acid-binding lectins during host cell invasion by apicomplexan parasites. J. Biol. Chem. 285, 2064–2076.

77. Marchant, J., Cowper, B., Liu, Y., Lai, L., Pinzan, C., Marq, J.B., Friedrich, N., Sawmynaden, K., Liew, L., Chai, W., et al. (2012). Galactose recognition by the apicomplexan parasite Toxoplasma gondii. J. Biol. Chem. 287, 16720–16733.

78. Patnaik, S.K., and Stanley, P. (2006). Lectin-resistant CHO glycosylation mutants. Methods Enzymol. 416, 159–182.

79. Esko, J.D., Wandall, H.H., and Stanley, P. (2022). Glycosylation mutants of cultured mammalian cells. In Essentials of Glycobiology (Cold Spring Harbor Laboratory Press), pp. 663–674.

80. Pinzan, C.F., Sardinha-Silva, A., Almeida, F., Lai, L., Lopes, C.D., Lourenço, E.V., Panunto-Castelo, A., Matthews, S., and Roque-Barreira, M.C. (2015). Vaccination with recombinant microneme proteins confers protection against experimental toxoplasmosis in mice. PLoS One 10, e0143087.

81. Lourenço, E.V., Pereira, S.R., Faça, V.M., Coelho-Castelo, A.A., Mineo, J.R., Roque-Barreira, M.C., Greene, L.J., and Panunto-Castelo, A. (2001). Toxoplasma gondii micronemal protein MIC1 is a lactose-binding lectin. Glycobiology 11, 541–547.

82. Sardinha-Silva, A., Mendonça-Natividade, F.C., Pinzan, C.F., Lopes, C.D., Costa, D.L., Jacot, D., Fernandes, F.F., Zorzetto-Fernandes, A.L.V., Gay, N.J., Sher, A., et al. (2019). The lectin-specific activity of Toxoplasma gondii microneme proteins 1 and 4 binds Toll-like receptor 2 and 4 N-glycans to regulate innate immune priming. PLoS Pathog 15, e1007871.

83. Reiss, M., Viebig, N., Brecht, S., Fourmaux, M.N., Soete, M., Di Cristina, M., Dubremetz, J.F., and Soldati, D. (2001). Identification and characterization of an escorter for two secretory adhesins in Toxoplasma gondii. J. Cell Biol. 152, 563–578.

84. Zhu, J., Wang, Y., Cao, Y., Shen, J., and Yu, L. (2021). Diverse roles of TgMIC1/4/6 in the Toxoplasma infection. Front. Microbiol. 12, 666506.

85. Possenti, A., Di Cristina, M., Nicastro, C., Lunghi, M., Messina, V., Piro, F., Tramontana, L., Cherchi, S., Falchi, M., Bertuccini, L., et al. (2022). Functional Characterization of the Thrombospondin-Related Paralogous Proteins Rhoptry Discharge Factors 1 and 2 Unveils Phenotypic Plasticity in Rhoptry Exocytosis. Front Microbiol 13, 899243.

86. Saouros, S., Edwards-Jones, B., Reiss, M., Sawmynaden, K., Cota, E., Simpson, P., Dowse, T.J., Jäkle, U., Ramboarina, S., Shivarattan, T., et al. (2005). A novel galectin-like domain from Toxoplasma gondii micronemal protein 1 assists the folding, assembly, and transport of a cell adhesion complex. J. Biol. Chem. 280, 38583–38591.

87. Huet, G., Hennebicq-Reig, S., de Bolos, C., Ulloa, F., Lesuffleur, T., Barbat, A., Carrière, V., Kim, I., Real, F.X., Delannoy, P., et al. (1998). GalNAc-alpha-O-benzyl inhibits NeuAcalpha2-3 glycosylation and blocks the intracellular transport of apical glycoproteins and mucus in differentiated HT-29 cells. J Cell Biol 141, 1311–1322.

88. Ulloa, F., Franci, C., and Real, F.X. (2000). GalNAc-alpha - O-benzyl inhibits sialylation of de Novo synthesized apical but not basolateral sialoglycoproteins and blocks lysosomal enzyme processing in a post-trans-Golgi network compartment. J. Biol. Chem. 275, 18785– 18793.

89. Lakshminarayan, R., Wunder, C., Becken, U., Howes, M.T., Benzing, C., Arumugam, S., Sales, S., Ariotti, N., Chambon, V., Lamaze, C., et al. (2014). Galectin-3 drives glycosphingolipid-dependent biogenesis of clathrin-independent carriers. Nat. Cell Biol. 16, 595–606.

90. Johannes, L., Parton, R.G., Bassereau, P., and Mayor, S. (2015). Building endocytic pits without clathrin. Nat. Rev. Mol. Cell Biol. 16, 311–321.

91. Ayyar, B.V., Ettayebi, K., Salmen, W., Karandikar, U.C., Neill, F.H., Tenge, V.R., Crawford, S.E., Bieberich, E., Prasad, B.V.V., Atmar, R.L., et al. (2023). CLIC and membrane wound repair pathways enable pandemic norovirus entry and infection. Nat. Commun. 14, 1148.

92. Liu, F.-T., and Stowell, S.R. (2023). The role of galectins in immunity and infection. Nat. Rev. Immunol. 23, 479–494.

93. Gagneux, P., Panin, V., Hennet, T., Aebi, M., and Varki, A. (2022). Evolution of glycan diversity. In Essentials of Glycobiology (Cold Spring Harbor Laboratory Press), pp. 265–278.

94. Viswanathan, K., Chandrasekaran, A., Srinivasan, A., Raman, R., Sasisekharan, V., and Sasisekharan, R. (2010). Glycans as receptors for influenza pathogenesis. Glycoconj. J. 27, 561–570.

95. Air, G.M. (2014). Influenza virus-glycan interactions. Curr. Opin. Virol. 7, 128–133.

96. Dessenne, C., Mariller, C., Vidal, O., Huvent, I., Guerardel, Y., Elass-Rochard, E., and Rossez, Y. (2025). Glycan-mediated adhesion mechanisms in antibiotic-resistant bacteria. BBA Adv. 7, 100156.

97. Lujan, A.L., Croci, D.O., Gambarte Tudela, J.A., Losinno, A.D., Cagnoni, A.J., Mariño, K.V., Damiani, M.T., and Rabinovich, G.A. (2018). Glycosylation-dependent galectin-receptor interactions promote Chlamydia trachomatis infection. Proc. Natl. Acad. Sci. U. S. A. 115, E6000–E6009.

98. Ashwood, C., and Cummings, R.D. (2025). N-glycopedia: Libraries for native N-glycan structural analysis. bioRxivorg, 2025.06.09.658590. 10.1101/2025.06.09.658590.

99. Otaki, M., Hirane, N., Natsume-Kitatani, Y., Nogami Itoh, M., Shindo, M., Kurebayashi, Y., and Nishimura, S.-I. (2022). Mouse tissue glycome atlas 2022 highlights inter-organ variation in major N-glycan profiles. Sci. Rep. 12, 17804.

100. Sauer, M.M., Jakob, R.P., Luber, T., Canonica, F., Navarra, G., Ernst, B., Unverzagt, C., Maier, T., and Glockshuber, R. (2019). Binding of the bacterial adhesin FimH to its natural, multivalent high-mannose type glycan targets. J. Am. Chem. Soc. 141, 936–944.

101. Ashwood, H.E., Ashwood, C., Schmidt, A.P., Gundry, R.L., Hoffmeister, K.M., and Anani, W.Q. (2021). Characterization and statistical modeling of glycosylation changes in sickle cell disease. Blood Adv. 5, 1463–1473.

102. Hirabayashi, J., Hashidate, T., Arata, Y., Nishi, N., Nakamura, T., Hirashima, M., Urashima, T., Oka, T., Futai, M., Muller, W.E.G., et al. (2002). Oligosaccharide specificity of galectins: a search by frontal affinity chromatography. Biochim. Biophys. Acta 1572, 232–254.

103. Zhuo, Y., and Bellis, S.L. (2011). Emerging role of alpha2,6-sialic acid as a negative regulator of galectin binding and function. J. Biol. Chem. 286, 5935–5941.

104. Zhuo, Y., Chammas, R., and Bellis, S.L. (2008). Sialylation of beta1 integrins blocks cell adhesion to galectin-3 and protects cells against galectin-3-induced apoptosis. J. Biol. Chem. 283, 22177–22185.

105. Sawmynaden, K., Saouros, S., Friedrich, N., Marchant, J., Simpson, P., Bleijlevens, B., Blackman, M.J., Soldati-Favre, D., and Matthews, S. (2008). Structural insights into microneme protein assembly reveal a new mode of EGF domain recognition. EMBO Rep. 9, 1149–1155.

106. Xie, Z., Angioletti-Uberti, S., Dobnikar, J., Frenkel, D., and Curk, T. (2025). Receptor clustering tunes and sharpens the selectivity of multivalent binding. Proc. Natl. Acad. Sci. U. S. A. 122, e2417159122.

107. Cerutti, A., Blanchard, N., and Besteiro, S. (2020). The bradyzoite: A key developmental stage for the persistence and pathogenesis of toxoplasmosis. Pathogens 9, 234.

108. Matta, S.K., Rinkenberger, N., Dunay, I.R., and Sibley, L.D. (2021). Toxoplasma gondii infection and its implications within the central nervous system. Nat. Rev. Microbiol. 19, 467–480.

109. Ahiya, A.I., Bhatnagar, S., Morrisey, J.M., Beck, J.R., and Vaidya, A.B. (2022). Dramatic consequences of reducing erythrocyte membrane cholesterol on Plasmodium falciparum. Microbiol. Spectr. 10, e0015822.

110. Maier, A.G., and van Ooij, C. (2022). The role of cholesterol in invasion and growth of malaria parasites. Front. Cell. Infect. Microbiol. 12, 984049.

111. Zidovetzki, R., and Levitan, I. (2007). Use of cyclodextrins to manipulate plasma membrane cholesterol content: evidence, misconceptions and control strategies. Biochim. Biophys. Acta 1768, 1311–1324.

112. Yang, J., Guo, F., Chin, H.S., Chen, G.B., Ang, C.H., Lin, Q., Hong, W., and Fu, N.Y. (2023). Sequential genome-wide CRISPR-Cas9 screens identify genes regulating cell-surface expression of tetraspanins. Cell Rep. 42, 112065.

113. Candor, K., Ding, L., Balchand, S., Hammonds, J.E., and Spearman, P. (2025). The CLIC/GEEC pathway regulates particle uptake and formation of the virus-containing compartment (VCC) in HIV-1-infected macrophages. PLoS Pathog. 21, e1012564.

114. Charron, A.J., and Sibley, L.D. (2004). Molecular partitioning during host cell penetration by Toxoplasma gondii. Traffic 5, 855–867.

115. Shen, B., and Sibley, L.D. (2012). The moving junction, a key portal to host cell invasion by apicomplexan parasites. Curr. Opin. Microbiol. 15, 449–455.

116. Shewell, L.K., Day, C.J., Jen, F.E.-C., Haselhorst, T., Atack, J.M., Reijneveld, J.F., Everest-Dass, A., James, D.B.A., Boguslawski, K.M., Brouwer, S., et al. (2020). All major cholesterol-dependent cytolysins use glycans as cellular receptors. Sci. Adv. 6, eaaz4926.

117. Ramakrishnan, C., Maier, S., Walker, R.A., Rehrauer, H., Joekel, D.E., Winiger, R.R., Basso, W.U., Grigg, M.E., Hehl, A.B., Deplazes, P., et al. (2019). An experimental genetically attenuated live vaccine to prevent transmission of Toxoplasma gondii by cats. Sci. Rep. 9, 1474.

118. Waldman, B.S., Schwarz, D., Wadsworth, M.H., Saeij, J.P., Shalek, A.K., and Lourido, S. (2020). Identification of a Master Regulator of Differentiation in Toxoplasma. Cell 180, 359– 372.e16.

119. Dubey, J.P., Lindsay, D.S., and Speer, C.A. (1998). Structures of*Toxoplasma gondii*tachyzoites, bradyzoites, and sporozoites and biology and development of tissue cysts. Clin. Microbiol. Rev. 11, 267–299.

120. Kent, R.S., and Ward, G.E. (2025). Motility-dependent processes in Toxoplasma gondii tachyzoites and bradyzoites: same same but different. mSphere 10, e0085524.

121. Lourenço, E.V., Bernardes, E.S., Silva, N.M., Mineo, J.R., Panunto-Castelo, A., and Roque-Barreira, M.-C. (2006). Immunization with MIC1 and MIC4 induces protective immunity against Toxoplasma gondii. Microbes Infect. 8, 1244–1251.

122. Ismael, A.B., Dimier-Poisson, I., Lebrun, M., Dubremetz, J.-F., Bout, D., and Mevelec, M.-N. (2006). Mic1-3 knockout of Toxoplasma gondii is a successful vaccine against chronic and congenital toxoplasmosis in mice. J. Infect. Dis. 194, 1176–1183.

123. Bojar, D., Meche, L., Meng, G., Eng, W., Smith, D.F., Cummings, R.D., and Mahal, L.K. (2022). A useful guide to lectin binding: Machine-learning directed annotation of 57 unique lectin specificities. ACS Chem. Biol. 17, 2993–3012.

124. Markus, B.M., Bell, G.W., Lorenzi, H.A., and Lourido, S. (2019). Optimizing Systems for Cas9 Expression in Toxoplasma gondii. Preprint, 10.1128/msphere.00386-19 https://doi.org/10.1128/msphere.00386-19.

125. Markus, B.M., Waldman, B.S., Lorenzi, H.A., and Lourido, S. (2020). High-Resolution Mapping of Transcription Initiation in the Asexual Stages of. Front Cell Infect Microbiol 10, 617998.

126. Smith, T.A., Lopez-Perez, G.S., Herneisen, A.L., Shortt, E., and Lourido, S. (2022). Screening the Toxoplasma kinome with high-throughput tagging identifies a regulator of invasion and egress. Nat. Microbiol. 7, 868–881.

127. Achbarou, A., Mercereau-Puijalon, O., Autheman, J.M., Fortier, B., Camus, D., and Dubremetz, J.F. (1991). Characterization of microneme proteins of Toxoplasma gondii. Mol. Biochem. Parasitol. 47, 223–233.

128. Schindelin, J., Arganda-Carreras, I., Frise, E., Kaynig, V., Longair, M., Pietzsch, T., Preibisch, S., Rueden, C., Saalfeld, S., Schmid, B., et al. (2012). Fiji: an open-source platform for biological-image analysis. Nat. Methods 9, 676–682.

129. Brown, K.M., Long, S., and Sibley, L.D. (2017). Plasma Membrane Association by N-Acylation Governs PKG Function in Toxoplasma gondii. MBio 8. 10.1128/mBio.00375-17.

130. Long, S., Brown, K.M., Drewry, L.L., Anthony, B., Phan, I.Q.H., and Sibley, L.D. (2017). Calmodulin-like proteins localized to the conoid regulate motility and cell invasion by Toxoplasma gondii. PLoS Pathog. 13, e1006379.

131. Wang, B., Wang, M., Zhang, W., Xiao, T., Chen, C.-H., Wu, A., Wu, F., Traugh, N., Wang, X., Li, Z., et al. (2019). Integrative analysis of pooled CRISPR genetic screens using MAGeCKFlute. Nat Protoc 14, 756–780.

132. Korotkevich, G., Sukhov, V., Budin, N., Shpak, B., Artyomov, M.N., and Sergushichev, A. (2016). Fast gene set enrichment analysis. bioRxiv, 060012. 10.1101/060012.

133. Burg, J.L., Perelman, D., Kasper, L.H., Ware, P.L., and Boothroyd, J.C. (1988). Molecular analysis of the gene encoding the major surface antigen of Toxoplasma gondii. J Immunol 141, 3584–3591.

134. Hughes, C.S., Moggridge, S., Müller, T., Sorensen, P.H., Morin, G.B., and Krijgsveld, J. (2019). Single-pot, solid-phase-enhanced sample preparation for proteomics experiments. Nat. Protoc. 14, 68–85.

